# The mechanism of analgesia in Na_V_1.7 null mutants

**DOI:** 10.1101/2020.06.01.127183

**Authors:** Donald Iain MacDonald, Shafaq Sikandar, Jan Weiss, Martina Pyrski, Ana P. Luiz, Queensta Millet, Edward C. Emery, Flavia Mancini, Gian D. Iannetti, Sascha R.A. Alles, Jing Zhao, James J Cox, Robert M. Brownstone, Frank Zufall, John N. Wood

**Affiliations:** Molecular Nociception Group, Wolfson Institute for Biomedical Research, University College London, Gower Street, WC1E 6BT, London; Centre for Experimental Medicine & Rheumatology, Queen Mary University of London, Charterhouse Square, EC1M 6BQ, London; Department of Physiology, University of Saarland School of Medicine, 66421 Homburg, Germany; Department of Neuroscience, Physiology and Pharmacology, University College London, Gower Street, WC1E 6BT, London; UCL Queen Square Institute of Neurology, WC1N 3BG, London

**Keywords:** pain, analgesia, sodium channels, Na_V_1.7, human genetics, endogenous opioids, neurotransmitter release

## Abstract

Deletion of *SCN9A* encoding the voltage-gated sodium channel Na_V_1.7 in humans leads to profound pain insensitivity and anosmia. Conditional deletion of Na_V_1.7 in sensory neurons of mice also abolishes pain suggesting the locus of analgesia is the nociceptor. Here we demonstrate that Na_V_1.7 knockout mice have essentially normal nociceptor activity using *in vivo* calcium imaging and extracellular recording. However, glutamate and substance P release from nociceptor central terminals in the spinal cord is greatly reduced by an opioid-dependent mechanism. Analgesia is also substantially reversed by central but not peripheral application of opioid antagonists. In contrast, the lack of neurotransmitter release from olfactory sensory neurons is opioid-independent. Male and female humans with Na_V_1.7 null mutations show naloxone reversible analgesia. Thus opioid-dependent inhibition of neurotransmitter release is the principal mechanism of Na_V_1.7 null analgesia in mice and humans.

## Introduction

Chronic pain afflicts a fifth of the population, but effective analgesics are few.^1^ We urgently need new molecular targets to develop improved painkillers. One strategy is to identify genes involved in rare human monogenic pain disorders. Loss-of-function mutations in the gene *SCN9A* encoding voltage-gated sodium channel Na_V_1.7 lead to congenital insensitivity to pain (CIP) and anosmia, but innocuous sensation remains intact.^2,3^ Gain-of-function mutations in *SCN9A* are associated with ongoing pain.^4–6^ Given the enriched expression of Na_V_1.7 in nociceptors, these discoveries point to a key role for Na_V_1.7 in controlling nociception in humans.^7^ Because Na_V_1.7 null individuals are wholly pain free, this channel represents a promising, human-validated drug target for pain relief.

Paradoxically, pharmacological block of the channel does not, in general, recapitulate the analgesia associated with functional deletion of the *SCN9A* gene.^8^ It has been assumed that Na_V_1.7 plays a key role in action potential initiation in the peripheral nerve endings of nociceptors, but neurogenic inflammation dependent on action potential propagation is not compromised in mouse or human Na_V_1.7 nulls.^9–11^ Loss of Na_V_1.7 also leads to enhanced endogenous opioid signalling and opioid receptor function in sensory neurons, and the opioid antagonist naloxone substantially reverses analgesia associated with channel deletion.^12–14^ Conditional knockout of *Scn9a* in the peripheral sensory neurons of mice reproduces the pain insensitivity of Na_V_1.7 null humans, affirming the nociceptor as the locus of analgesia. These animals show profound behavioural deficits in thermal, mechanical, inflammatory and some forms of neuropathic pain.^15,16^ To determine the mechanism of Na_V_1.7 null analgesia, we used complementary optical, electrophysiological and pharmacological methods to study nociceptor function *in vivo* in mice and humans lacking Na_V_1.7.

## Results

### Deletion of Na_V_1.7 in sensory neurons decreases pain sensitivity without silencing peripheral nociceptors

We deleted *Scn9a* encoding Na_V_1.7 in peripheral sensory neurons of mice. Advillin-Cre or Wnt1-Cre was used to excise the floxed *Scn9a* allele, resulting in knockout of Na_V_1.7 restricted, respectively, to sensory neurons or to neural crest-derived neurons.^15^ We performed whole-cell voltage-clamp recordings of voltage-gated sodium currents in cultured sensory neurons from control animals homozygous for the floxed *Scn9a* allele, here called wild-type (WT). Application of the Na_V_1.7-specific antagonist PF-05089771 (PF-771, 100 nM for 5 minutes) showed that ∼50% of the peak sodium current in medium diameter nociceptor-like neurons can be attributed to Na_V_1.7 (Figure S1A).^17^ In contrast, PF-771 had no effect on voltage-gated sodium currents recorded in sensory neurons from either Advillin-Cre Na_V_1.7 knockout (KO^Adv^) or Wnt1-Cre Na_V_1.7 knockout (KO^Wnt^) mice, confirming functional loss of the channel (Figure S1B & S1C). In common with human Na_V_1.7 null individuals, both lines of conditional Na_V_1.7 KOs showed a profound analgesic phenotype characterized by increased withdrawal thresholds to noxious thermal and mechanical stimuli, but intact responses to innocuous cold and tactile stimuli (Figure S2A-D).

To test the hypothesis that loss of Na_V_1.7 silences nociceptors, we monitored the responses of peripheral sensory neurons to noxious stimulation using *in vivo* calcium imaging.^18^ We generated conditional Na_V_1.7 KO and WT mice on a Pirt-GCaMP3 background, where GCaMP3 is found in all peripheral sensory neurons.^19^ Using laser-scanning confocal microscopy, we imaged calcium signals in sensory neuron somata within the L4 dorsal root ganglia of live anesthetised animals (Figure 1A). Na_V_1.7-deficient sensory neurons readily responded to all noxious stimuli applied to the hindpaw (Figure 1B). We classified responding neurons into functionally-defined cell types. Every cell type was present in both WT and KO animals, but differences were apparent in the distribution of responses (Figure 1C). Fewer cells responded to noxious pinch in KO^Adv^ (44/197, 22%) and KO^Wnt^ (70/262, 27%) compared to WT (206/516, 40%). This could be linked to loss of excitability in some nociceptors observed in *in vitro* culture experiments.^20^ We observed a corresponding increase in the proportion of cold-sensing neurons in KO^Adv^ (73/197, 37%) and KO^Wnt^ (117/262, 45%) versus WT (132/516, 26%). Importantly, the relative number of cells responding to noxious heat was not markedly altered between WT (234/516, 45%), KO^Adv^ (95/197, 48%) and KO^Wnt^ (88/262, 34%) lines, despite behavioural insensitivity to heat (Figure S2C). We wondered whether polymodal nociceptors were silenced by loss of Na_V_1.7, here defined as pinch-sensitive cells that also respond to thermal stimuli. Polymodality did not differ between genotypes and was similar to previous reports, with 21% of WT, 25% of KO^Adv^ and 16% of KO^Wnt^ neurons categorized as polymodal (Figure 1D).^18,21–23^ Lastly, we measured the peak calcium signal (ΔF/F_0_) as a surrogate measure of single neuron excitability. When we quantified this for each stimulus type, there was no effect of Na_V_1.7 deletion on the maximum calcium responses (Figure 1E). Overall, although the distribution of cold and mechanical responses was altered, we found little evidence using calcium imaging of decreased nociceptor excitability in animals lacking Na_V_1.7, with no change in peak response magnitude to any stimulus, prevalence of polymodality or number of noxious heat responses. Broadly similar results were obtained by calcium imaging of WT and KO^Adv^ mice virally expressing GCaMP6f (Figure S3A-D).

**Figure 1.**
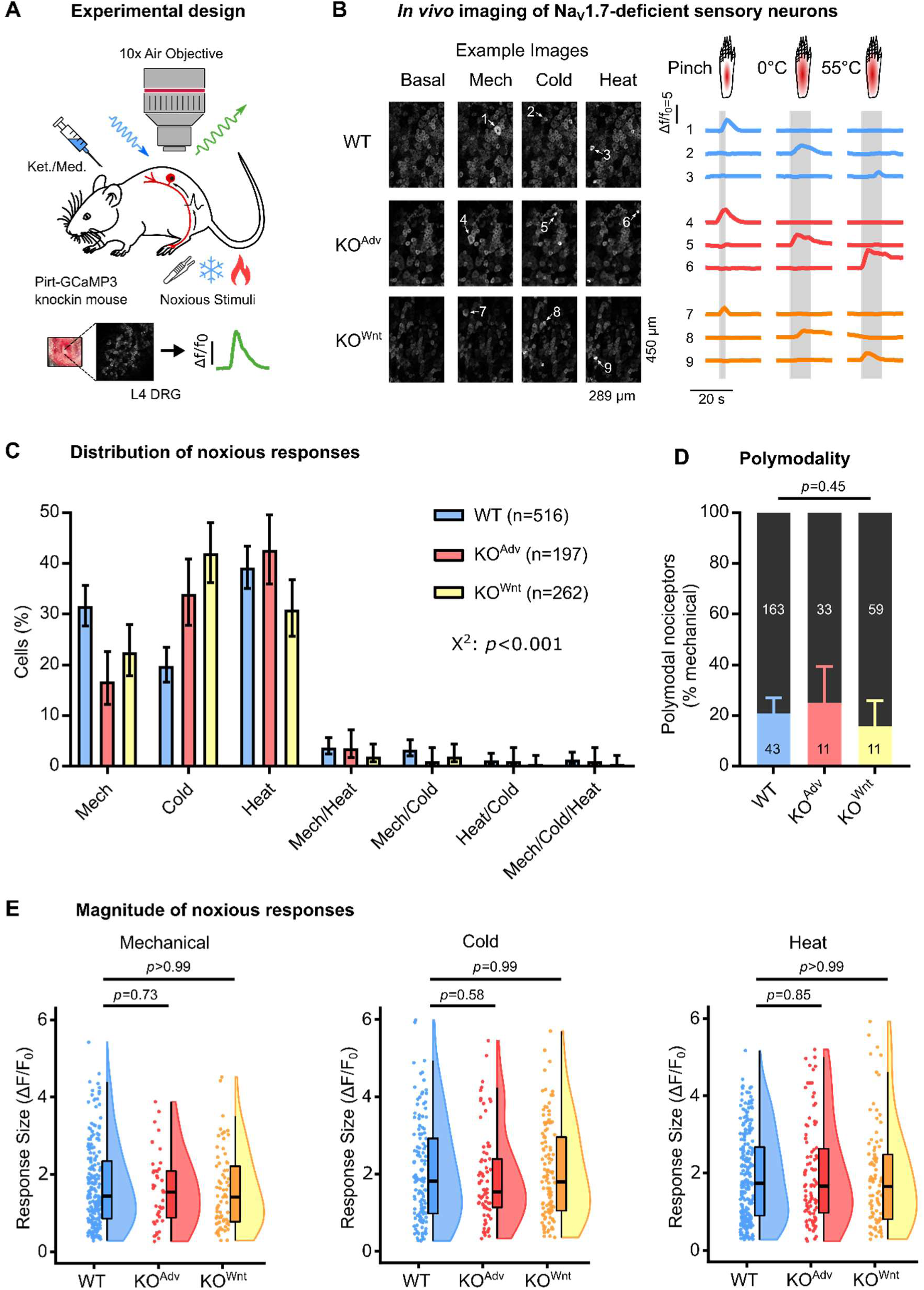
Na_V_1.7-deficient sensory neurons respond to noxious stimuli at the level of the soma, *in vivo*. (A) Schematic of *in vivo* dorsal root ganglion imaging setup. (B) Example images and traces showing sensory neurons respond to noxious mechanical and thermal stimuli in WT and both Na_V_1.7 KO mouse lines. Each numbered trace corresponds to one cell. The data in this figure were obtained from 19 WT, 8 KO^Adv^ and 11 KO^Wnt^ animals. (C) Bar plot summarizing the distribution of all sensory neurons that responded to different noxious stimuli in WT and Na_V_1.7 KO animals. The error bars represent 95% confidence intervals and proportions were compared using Chi-Square (χ^2^) test. n=516 cells from WT (blue), n=187 cells from KO^Adv^ (red) and n=262 cells from KO^Wnt^ (yellow). (D) Bar plot showing similar prevalence of polymodal nociceptors in WT and Na_V_1.7 KO mice. Polymodal nociceptors are defined as pinch-sensitive neurons that respond to any noxious thermal stimulus (colour) and are expressed as a fraction of mechanically-sensitive cells (black). The error bars represent 95% confidence intervals and proportions were compared using the Chi-Square test. n=206 cells from WT, n=44 cells from KO^Adv^ and n=70 cells from KO^Wnt^. (E-F) Raincloud plots showing similar peak calcium responses (ΔF/F_0_) evoked by different noxious stimuli for WT and Na_V_1.7 KO lines. Mean response magnitude of KO lines was compared to WT control using One-Way ANOVA followed by post-hoc Dunnett’s test. Mechanical: n=206 cells from WT, n=44 cells from KO^Adv^ and n=70 cells from KO^Wnt^. Cold: n=132 cells from WT, n=73 cells from KO^Adv^ and n=117 cells from KO^Wnt^. Heat: n=234 cells from WT, n=95 cells from KO^Adv^ and n=88 cells from KO^Wnt^.

While calcium imaging is ideally suited to monitoring population activity, we cannot directly measure action potential firing. We reasoned that this method may not be sensitive to subtler effects of Na_V_1.7 deletion on excitability. We therefore performed multi-unit extracellular recordings from dorsal root ganglia of live Na_V_1.7 KO and WT animals. For these experiments, we pooled data obtained from both KO^Adv^ and KO^Wnt^ lines, quantifying the number of action potentials fired in 10 seconds by polymodal afferents to peripheral stimuli of various modalities (Figure 2A). Firing evoked by noxious mechanical prodding was unchanged after deletion of Na_V_1.7 (Figure 2B). Intriguingly, although firing in response to most von Frey stimuli was normal, suprathreshold responses to 8, 15 and 26 g gram hairs showed a small reduction (Figure 2C). This is in spite of the normal hind-paw von Frey thresholds of Na_V_1.7 KO mice (Figure S2B). As expected, innocuous brush stimuli evoked action potentials equally well in WT and KO animals. There was no appreciable change in the number of spikes triggered by ice water (Figure 2E) or noxious heat stimuli (Figure 2F). Taken together, these data show that the diminished sensitivity of Na_V_1.7 KO mice to noxious heat or mechanical stimuli cannot be explained by decreased peripheral nociceptor excitability.

**Figure 2.**
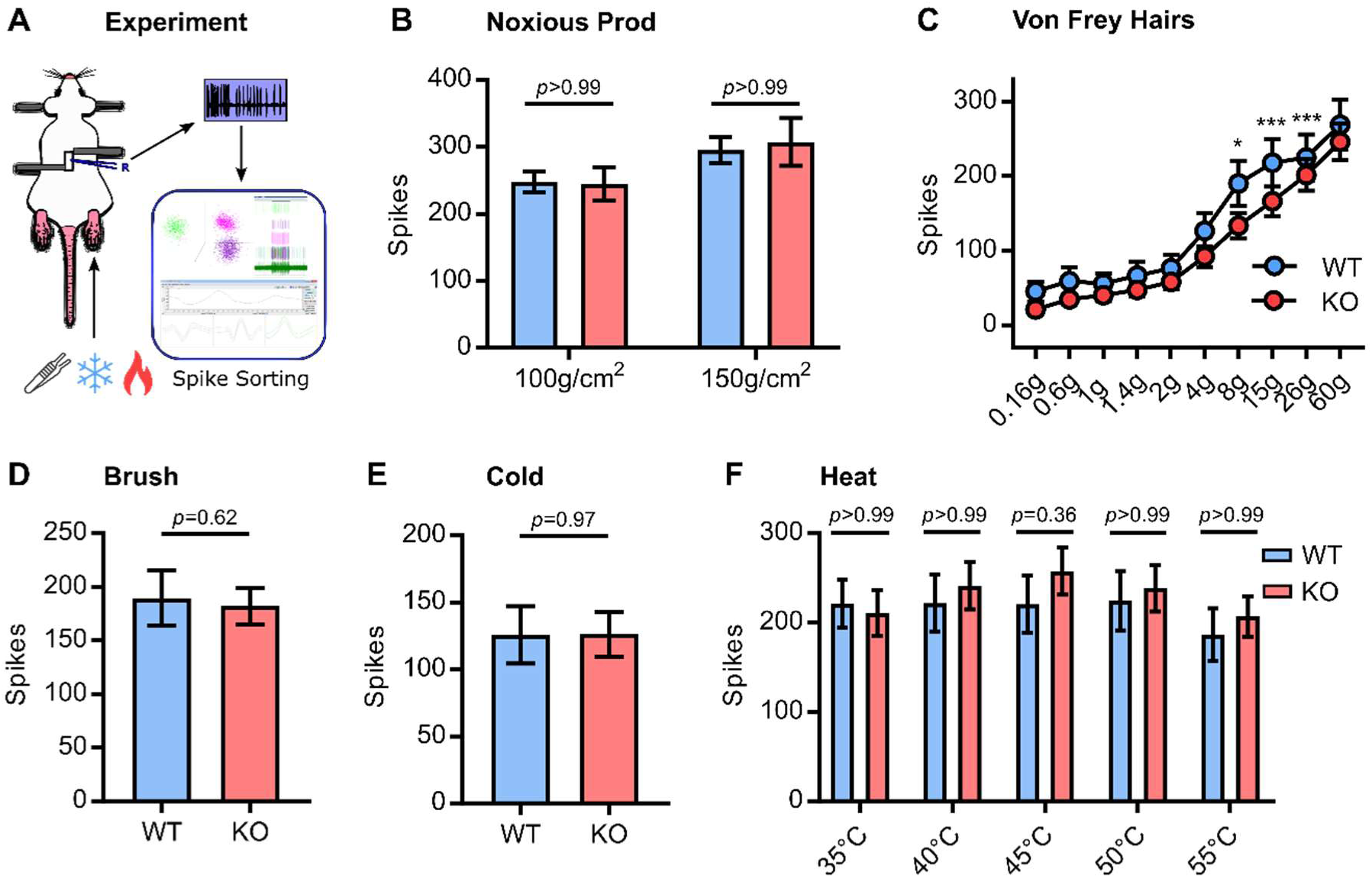
Excitability of Na_V_1.7-deficient sensory neurons, *in vivo*. (A) Schematic of *in vivo* dorsal root ganglion extracellular recording setup. (B) Quantification of spikes fired in response to noxious prodding in WT (blue) and Na_V_1.7 KO (red). Data from both Advillin-Cre and Wnt1-Cre Na_V_1.7 KO mice were pooled for these experiments. (C) Quantification of spikes fired in response to von Frey hair stimulation. For 8g, *p*=0.014. For 15g, *p*<0.001. For 26g, *p*<0.001. (D) Quantification of spikes fired in response to brushing. (E) Quantification of spikes fired in response to cooling. (F) Quantification of spikes fired in response to heating. For (B), (C), and (F) mean number of spikes fired in 10 s were compared using repeated measures Two-Way ANOVA followed by post-hoc Bonferroni test. For (D) and (E), means were compared using an unpaired t test. Error bars represent 95% confidence interval around the mean. n=90 cells from 10 WT animals and n=146 cells from 13 KO^Adv/Wnt^ animals.

Of particular clinical interest is the fact both mice and humans lacking Na_V_1.7 show deficient inflammatory pain sensitization. Prostaglandin E2 (PGE2) is an important inflammatory mediator.^18^ To investigate the mechanism by which Na_V_1.7 deletion impedes pain sensitization, we tested the effect of intraplantar injection of PGE2 (500 µM for 10 minutes) on whole-animal behavioural responses and sensory neuron calcium responses to noxious heat stimuli, *in vivo* (Figure 3A). Both lines of conditional Na_V_1.7 KO mice failed to develop heat hyperalgesia (Figure 3B). In contrast, intraplantar injection of PGE2 unmasked silent nociceptors and increased heat responses in both WT and KO^Adv^ animals expressing GCaMP6f, indicating this form of peripheral sensitization is intact in mice lacking Na_V_1.7 (Figure 3C). We also observed robust sensitizing effects of PGE2 in KO^Wnt^ animals expressing GCaMP3, although the number of silent nociceptors activated was slightly less than WT (Figure 3D). Na_V_1.7 deletion therefore impairs thermal hyperalgesia but paradoxically does not abolish its physiological correlate: the peripheral sensitization of nociceptors.

**Figure 3.**
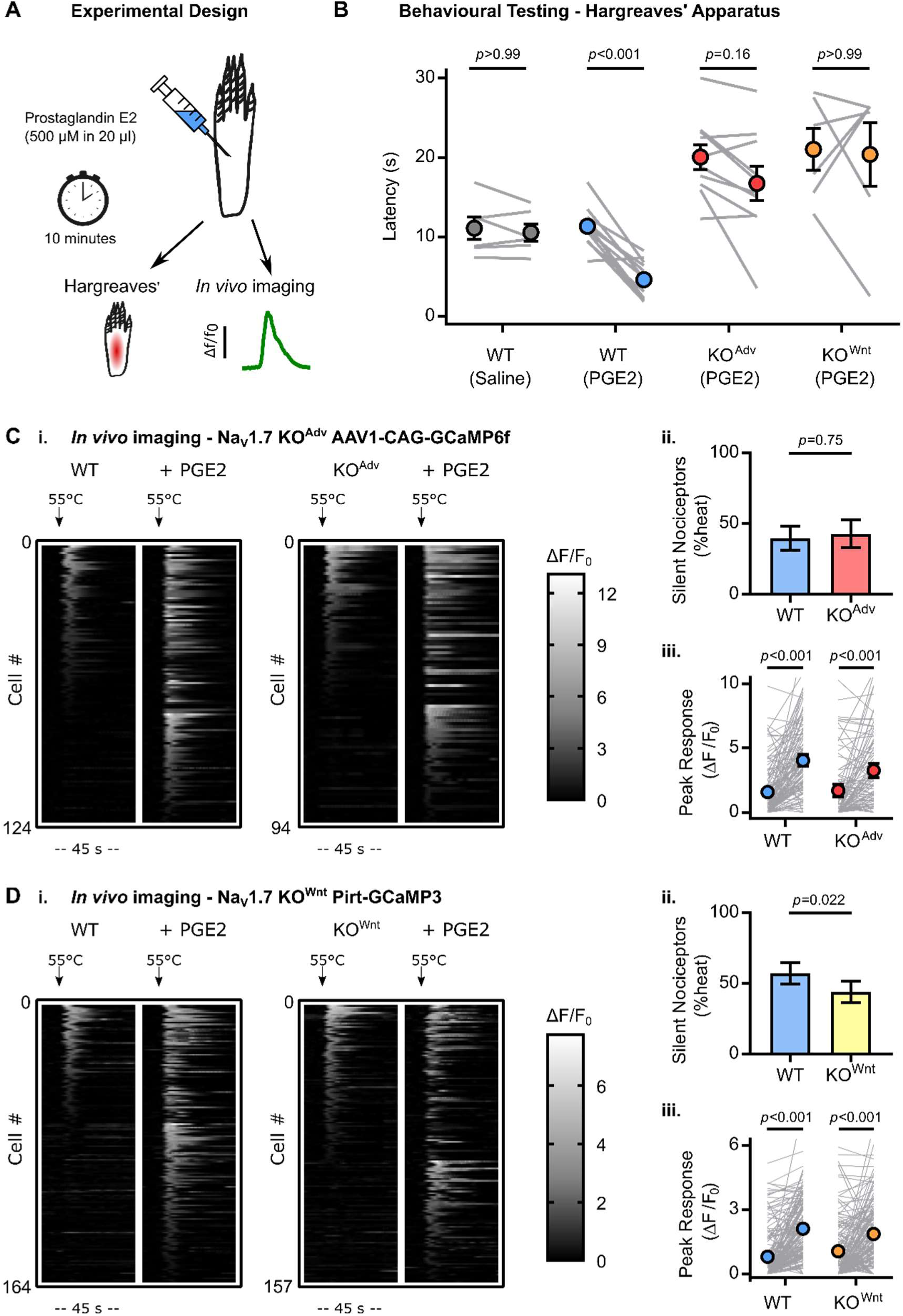
Na_V_1.7 deletion abolishes inflammatory pain without affecting peripheral sensitization. (A) Schematic illustrating induction of acute inflammatory pain using PGE2. (B) Behavioural assessment of the effect of PGE2 on Hargreaves’ withdrawal latencies in WT and Na_V_1.7 KO animals, showing KO mice do not develop heat hyperalgesia. The error bars represent standard error of the mean. Mean latencies before and after PGE2 were compared using repeated measures Two-Way ANOVA followed by post-hoc Sidak’s test. n=6 animals for WT vehicle, n=12 for WT PGE2, n=10 for KO^Adv^ PGE2, and n=6 for KO^Wnt^ PGE2. (C) Heat maps (i.) and quantification (ii. & iii.) showing unmasking of silent heat nociceptors by PGE2 in both WT and Na_V_1.7 KO^Adv^ animals virally transduced with GCaMP6f. n=124 cells from 4 WT animals and n=94 cells from 4 KO^Adv^. (D) Heat maps (i.) and quantification (ii. & iii.) showing unmasking of silent heat nociceptors by PGE2 in both WT and Na_V_1.7 KO^Wnt^ animals expressing GCaMP3. n=164 cells from 8 WT animals and n=157 cells from 11 KO^Wnt^ animals. For both (C) and (D), the effect of genotype on PGE2-induced unmasking of silent nociceptors was compared using the Chi-Square test with Yates’ Correction for proportions (i.) and repeated measures Two-Way ANOVA followed by post-hoc Sidak’s test for mean response size (ii.). Error bars represent 95% confidence intervals.

### Loss of Na_V_1.7 impairs synaptic transmission from nociceptors

The mechanism of Na_V_1.7 analgesia does not appear to arise from reduced peripheral excitability. We wondered whether the loss of function occurred at the nociceptor central terminal. To test this, we used the fluorescent glutamate sensor iGluSnFR to directly measure glutamate release from sensory afferent central terminals in spinal cord slices.^24^ First, we virally transfected cultured dorsal root ganglia neurons with iGluSnFR. iGluSnFR signal was present on the cell membrane and along neurites (Figure S4A). Bath application of a range of glutamate concentrations confirmed the sensitivity of the probe to extracellular glutamate concentrations within the physiological range (Figure S4B). We then virally transduced sensory neurons *in vivo* with iGluSnFR by intraperitoneal injection of AAV9-synapsin-iGluSnFR virus at P2 (Figure S4C). Spinal cord slices were prepared for 2-photon imaging experiments at P9-21. In the dorsal horn, we observed iGluSnFR fluorescence in the central processes of incoming afferents, but no spinal cord cell bodies were labelled (Figure S4D). In contrast, confocal imaging showed iGluSnFR was widely expressed in the cell bodies of sensory neurons in the dorsal root ganglia, *in vivo*, confirming the successful targeting of iGluSnFR to afferent neurons (Figure S4E).

To image synaptic transmission in the spinal cord, dorsal root stimulation was used to evoke neurotransmitter release from central afferent terminals expressing iGluSnFR in WT and KO^Adv^ mice (Figure 4A). We restricted our imaging of glutamate signals to lamina II in the dorsal horn of the spinal cord. Two-photon imaging at 10 Hz showed glutamate release was readily evoked across a range of single pulse stimulus intensities in dorsal horn of spinal cord slices, with iGluSnFR signals showing spatial localization to regions of interest (Figure 4B). Interestingly, the mean minimum stimulus current required to elicit release in slices from KO^Adv^ animals was 891 µA, threefold greater than the control wild type value of 279 µA (Figure 4Ci.). The EC_50_ current was also increased in the KO^Adv^ slices with a value of 366 µA compared to 181 µA in WT (Figure 4Cii). Even when highest intensity stimuli were used, the peak fluorescence change (ΔF/F_0_) was reduced in the KO^Adv^ slices (Figure 4Ciii). These data are consistent with a reduction in glutamate release at the central terminal of sensory neurons in Na_V_1.7 KO^Adv^ animals.

**Figure 4.**
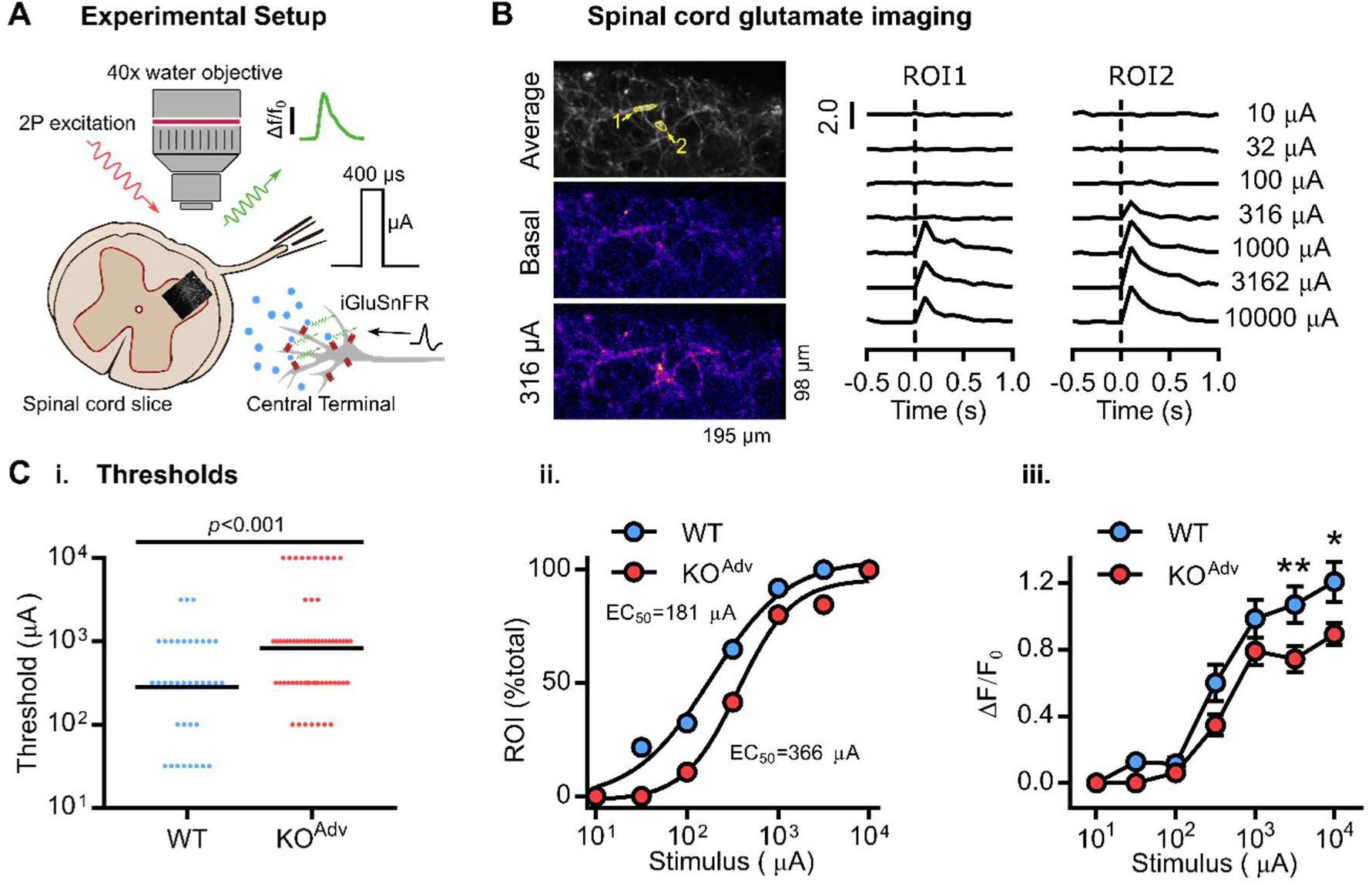
Decreased neurotransmitter release from the central terminals of Na_V_1.7-deficient sensory neurons. (A) Schematic illustrating two-photon imaging of iGluSnFR-expressing afferent terminals in the dorsal horn of spinal cord slices. Data were obtained from 4 WT and 6 KO^Adv^ animals. (B) Example images and traces of glutamate release from afferent terminals in response to single-pulse dorsal root stimulation in spinal cord slices obtained from a Na_V_1.7 KO^Adv^ mouse. ROI=region of interest. (C) Plots showing the threshold current required to evoke glutamate release is decreased in slices from KO^Adv^ mice. For (i.), the mean absolute threshold was compared between genotypes using an unpaired t test. For (ii.), WT: EC_50_=181 µA, r^2^=0.99 and KO^Adv^: EC_50_=366 µA, r^2^=0.99. For (iii.), mean evoked glutamate release (ΔF/F_0_) was compared between genotypes using repeated measures Two-Way ANOVA followed by post-hoc Sidak’s test. \*\**p*=0.007, \**p*=0.010. n=37 ROIs for WT. n=65 ROIs for KO^Adv^.

Next, we performed voltage-clamp recordings from lamina II neurons in spinal cord slices, which receive input from afferents expressing Na_V_1.7.^25^ When we examined spontaneous EPSCs, there was a reduction in frequency but not amplitude, consistent with impaired presynaptic release (Figure S5A-C). These presynaptic deficits may account for the heightened pain thresholds of mice lacking Na_V_1.7 and associated diminished nociceptive input to the CNS.^15^

### Opioid receptors are required for synaptic deficits in nociceptors but not olfactory sensory neurons lacking Na_V_1.7

We have previously shown that analgesia in mice and humans lacking Na_V_1.7 requires opioid receptors. This is due to both upregulation of preproenkephalin (PENK) and enhanced opioid receptor signalling caused by Na_V_1.7 deletion.^12–14^ Does increased opioid signalling account for the synaptic deficits we observe in Na_V_1.7 KOs? We first investigated the effect of systemic naloxone injection *in vivo* (2 mg/kg subcutaneously for 20 minutes) on peripheral excitability of nociceptors. To our surprise, *in vivo* imaging experiments revealed that naloxone increased the number of responding cells in both WT and KO^Adv^ mice, suggesting tonic endogenous opioid activity is present in our preparation (Figure 5A). Importantly, however, the effect of naloxone on the number and size of responses was comparable across genotypes. Corroborating this, *in vivo* extracellular recording of sensory neurons from WT and KO^Adv^ found no effect of naloxone on action potential firing in response to noxious heat, cold or mechanical stimuli (Figure 5B).

**Figure 5.**
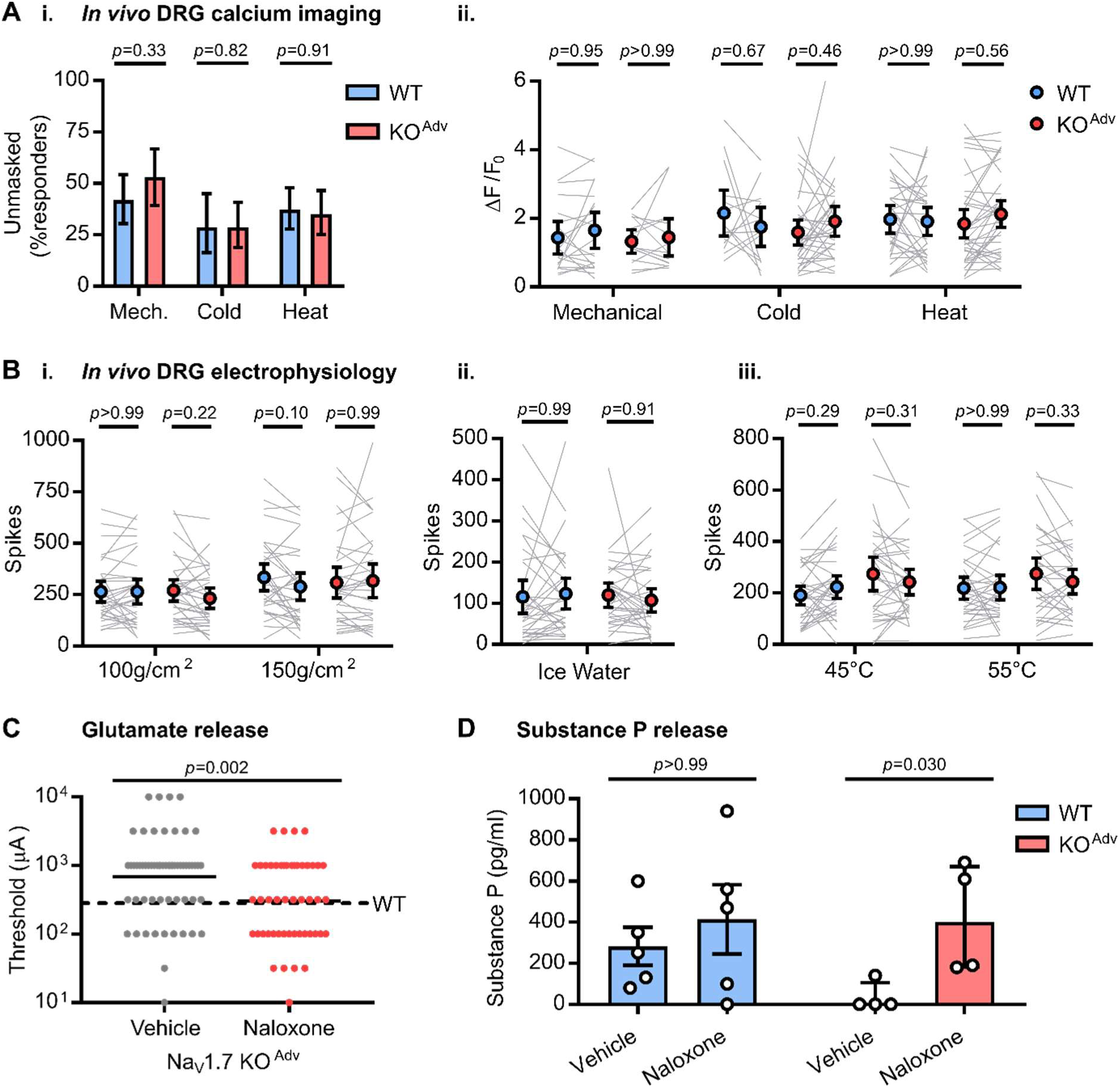
Opioid receptor blockade rescues impaired neurotransmission after Na_V_1.7 deletion but does not affect peripheral excitability. (A) *In vivo* imaging of sensory neuron activity before and after treatment with systemic naloxone (2 mg/kg subcutaneously for 20 minutes) in WT (blue) and Na_V_1.7 KO^Adv^ (red) mice. (i.) Naloxone unmasked previously silent neurons in both WT and KO^Adv^ mice. Proportions were compared using Chi square test with Yates’ correction. (ii.) Naloxone had no effect on the peak calcium responses evoked by noxious stimuli in either genotype. Mean peak calcium responses before and after naloxone were compared using repeated measures Two-Way ANOVA followed by post-hoc Sidak’s test. Data were obtained from 4 WT and 6 KO^Adv^ animals. Mechanical: n=62 cells from WT, n=47 cells from KO^Adv^. Cold: n=35 cells from WT, n=63 cells from KO^Adv^. Heat: n=86 cells from WT, n=74 cells from KO^Adv^. (B) *In vivo* extracellular recording of sensory neuron action potential firing before and after treatment with systemic naloxone (2 mg/kg subcutaneously for 20 minutes) in WT (blue) and Na_V_1.7 KO^Adv^ (red) mice. Naloxone had no effect on spiking evoked by noxious mechanical (i.), ice water (ii.) or heat (iii.) stimuli. Mean spikes fired before and after naloxone were compared using repeated measures Two-Way ANOVA followed by post-hoc Sidak’s test. n=33 from 6 WT animals, and n=33 from 7 KO^Adv^ animals. (C) *Ex vivo* iGluSnFR imaging of glutamate release from sensory neuron central terminals in dorsal horn of spinal cord slices from 6 KO^Adv^ animals treated with vehicle (grey) or 100 µM naloxone (red). Naloxone reduced mean threshold current required to elicit release to WT levels (dashed line). n=62 regions of interest for vehicle and n=50 for naloxone. (D) Measurement of substance P release from spinal cord slices obtained from 5 WT and 4 KO^Adv^ animals. Treatment with 1 mM naloxone rescued diminished substance P release in KO^Adv^ slices, but had no effect in WT. Data were compared using Kruskall Wallis test.

As naloxone did not markedly affect peripheral excitability, we examined the relationship between opioid receptors and synaptic dysfunction in Na_V_1.7 KOs. Spinal cord slices from Na_V_1.7 KO^Adv^ animals were treated with either vehicle or naloxone while measuring dorsal root stimulation-evoked glutamate release using iGluSnFR. Vehicle-treated KO^Adv^ slices showed a mean minimum stimulus current for eliciting release of 690 µA. This was reduced to 302 µA in naloxone-treated KO^Adv^ slices, comparable to the 279 µA threshold we previously observed in WT slices (Figure 5C). The EC_50_ current was 457 µA in the vehicle-treated group, and decreased to 173 µA after naloxone, essentially identical to the WT value of 181 µA. These findings indicate that opioid receptor blockade reverses changes in glutamate release associated with deletion of Na_V_1.7.

Next, we asked if the deficits in substance P release previously observed in KO^Adv^ mice are also dependent on opioid receptors.^15^ Electrical stimulation of hemisected spinal cord with dorsal roots attached was originally used to show a deficit in substance P release in KO^Adv^ mice. Here we used the potent pain-producing TRPV1 ligand capsaicin to selectively depolarise nociceptor terminals in spinal cord slices from WT and KO^Adv^ mice. Using a competitive binding immunoassay, we measured the amount of substance P release elicited by 2 µM capsaicin stimulation in both genotypes with and without 1 mM naloxone (Figure 5D). In KO^Adv^ slices, a mean of 35 ± 35 pg / ml of substance P was detected. Treatment with naloxone increased release to 418 ± 135 pg / ml. In WT slices, there was no significant difference between vehicle (282 ± 92 pg / ml) and naloxone conditions (414 ± 169 pg / ml). The diminution in substance P release from nociceptor central terminals lacking Na_V_1.7 is thus rescued by block of opioid receptors.

Mice lacking Na_V_1.7 in olfactory sensory neurons (Nav1.7 KO^OMP^) are totally anosmic due to a loss of transmitter release from olfactory nerve terminals.^3^ To investigate if these synaptic deficits are also dependent on opioid receptors, we recorded M/T cells in horizontal olfactory bulb slices of KO^OMP^ mice. As expected, electrical stimulation (1 ms, 100 V) of the olfactory nerve layer harbouring the olfactory sensory neuron axon terminals did not elicit a postsynaptic current in M/T cells of Nav1.7 KO^OMP^ (Figure 6A). To test if the opioid receptor antagonist naloxone could restore transmitter release at this synapse, we bath-applied 300 µM naloxone for at least 10 minutes and recorded olfactory nerve-evoked responses in 17 M/T cells (5 animals). Naloxone did not lead to an increase in EPSC amplitude (Figure 6B) (Nav1.7^control^ −14 ± 0.4 pA vs Nav1.7KO^OMP^ −13.9 ± 0.03 pA). To exclude the possibility that naloxone affects transmitter release in general at this synapse we additionally recorded M/T cells of control mice (Nav1.7^control^, 7 cells in 4 animals). Olfactory nerve stimulation caused activation of characteristic inward currents in M/T cells of control mice with amplitudes of −165 ± 20 pA. Application of naloxone to the slice had only minor effects on evoked EPSCs (−148 ± 37 pA) (Figure 6A&B). As further corroboration, animals were injected intraperitoneally 3 times (30 minute interval) with naloxone (2 mg/kg) or PBS and kept in the odour-rich environment of their homecage for the next 24 hours. Animals were then sacrificed, perfused and olfactory bulb coronal sections stained against tyrosine hydroxylase (TH) protein. TH expression in olfactory bulb juxtaglomerular neurons is a correlate of afferent synaptic input because it requires odour-stimulated glutamate release from OSN terminals.^3^ There was no obvious difference between PBS and naloxone treated animals in their amount of TH expression, with low to no expression of TH in KO^OMP^ mice, and high expression in control mice (Figure 6C). These results are consistent with the finding that, in Nav1.7^control^ mice, treatment with a cocktail of opioid receptor agonists did not affect EPSCs evoked by olfactory nerve stimulation (Figure 6D). TTX, on the other hand, completely abolished synaptic transmission from OSNs to M/T cells (Figure 6E).

**Figure 6.**
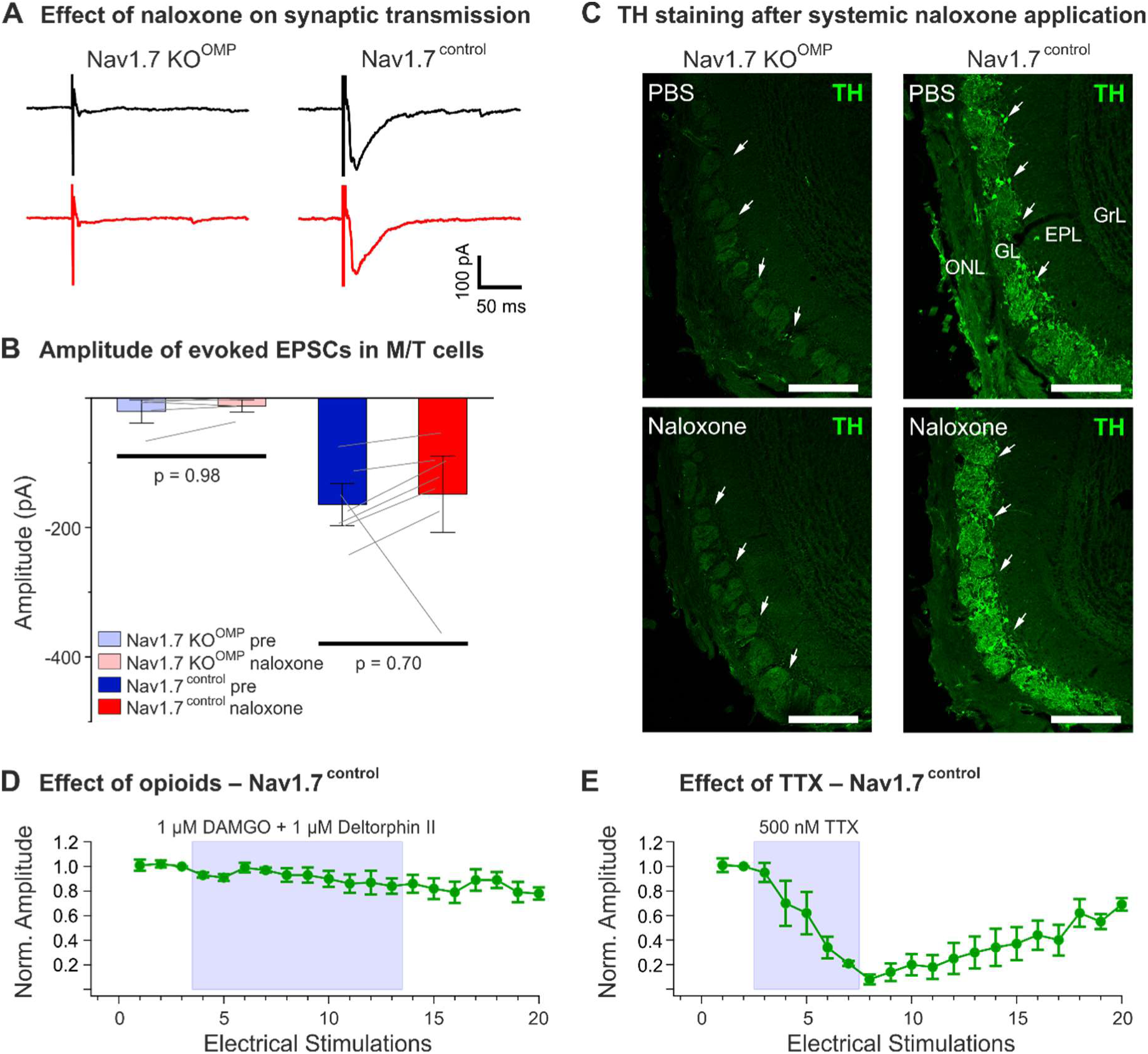
Opioid receptor block does not rescue synaptic transmission in mice lacking Na_V_1.7 in olfactory sensory neurons. (A) Loss of synaptic transmission onto M/T cells in olfactory bulb slices of Na_V_1.7 KO^OMP^ mice after olfactory sensory neuron nerve stimulation cannot be rescued by 300 µM naloxone (left, red trace). Additionally, 300 µM naloxone does not affect M/T EPSCs to presynaptic nerve stimulation in control mice (right, red trace). (B) Summary plot showing the morphine receptor antagonist naloxone (300 µM) does not affect EPSC amplitudes in M/T cells after presynaptic nerve stimulation; neither in Na_V_1.7 KO^OMP^ (KO^OMP^ pre and naloxone, n=17 from 5 animals) nor in control mice (pre and naloxone, n=7 from 4 animals). Error bars represent SEM. Means were compared using paired t test. (C) Confocal images of tyrosine hydroxylase (TH) immunostaining (green) in coronal cryosections of the major olfactory bulb (MOB) following systemic administration of PBS or naloxone of adult Na_V_1.7 KO^OMP^ (left) and control (right) mice. TH staining is absent in the glomerular layer (GL) of Na_V_1.7 KO^OMP^ mice independent of treatment (arrows), while the MOB of control mice shows robust TH labeling of neuronal processes and periglomerular cell somata (arrows). There is no difference in TH staining in control MOBs when comparing PBS vs naloxone administration. ONL (olfactory nerve layer), EPL (external plexifiorm layer), GrL (granule cell layer). Scale bars, 200 µm. (D) Time course showing treatment with 1 µM DAMGO and 1 µM Deltorphin II does not affect M/T cell EPSCs evoked by olfactory nerve stimulation (n=4). Normalized EPCS peak amplitudes are plotted as a function of the number of electrical ONL stimulation (1-min intervals). Error bars represent SEM. (E) Time course showing treatment with 500 nM TTX completely and reversibly abolishes M/T cell EPSCs evoked by ONL stimulation (n=4). Error bars represent standard error of the mean.

### Pain insensitivity of mice and humans lacking Na_V_1.7 depends on opioid signalling

Is the suppression of synaptic transmission by the opioid system required for analgesia? To test this in awake behaving animals, we selectively blocked opioid receptors in either the peripheral, central or both compartments of the nociceptor (Figure 7A). We measured withdrawal latencies to radiant heat stimuli as a read-out. To test if central opioid receptors are required, we injected naloxone (3 mM in 5 µl) by intrathecal injection into the lumbar spinal column. Centrally-administered naloxone was sufficient to reverse thermal hyposensitivity in Na_V_1.7 KO^Adv^ mice by 76%. There was no effect of intrathecal vehicle injection (Figure 7B). Naloxone methiodide is a peripherally-restricted naloxone analogue that does not cross the blood brain barrier and hence selectively blocks peripheral opioid receptors.^26^ Systemic administration of naloxone methiodide (2 mg/kg) had no impact on withdrawal latencies in either WT or KO^Adv^ mice, indicating peripheral opioid receptors are dispensable for analgesia linked to Na_V_1.7 loss-of-function (Figure 7C). In contrast, systemic injection of naloxone (2 mg/kg) caused a 71% reversal of analgesia in the KO^Adv^ group. Thus, central – but not peripheral – opioid receptors are essential for the maintenance of analgesia in mice lacking Na_V_1.7.

**Figure 7.**
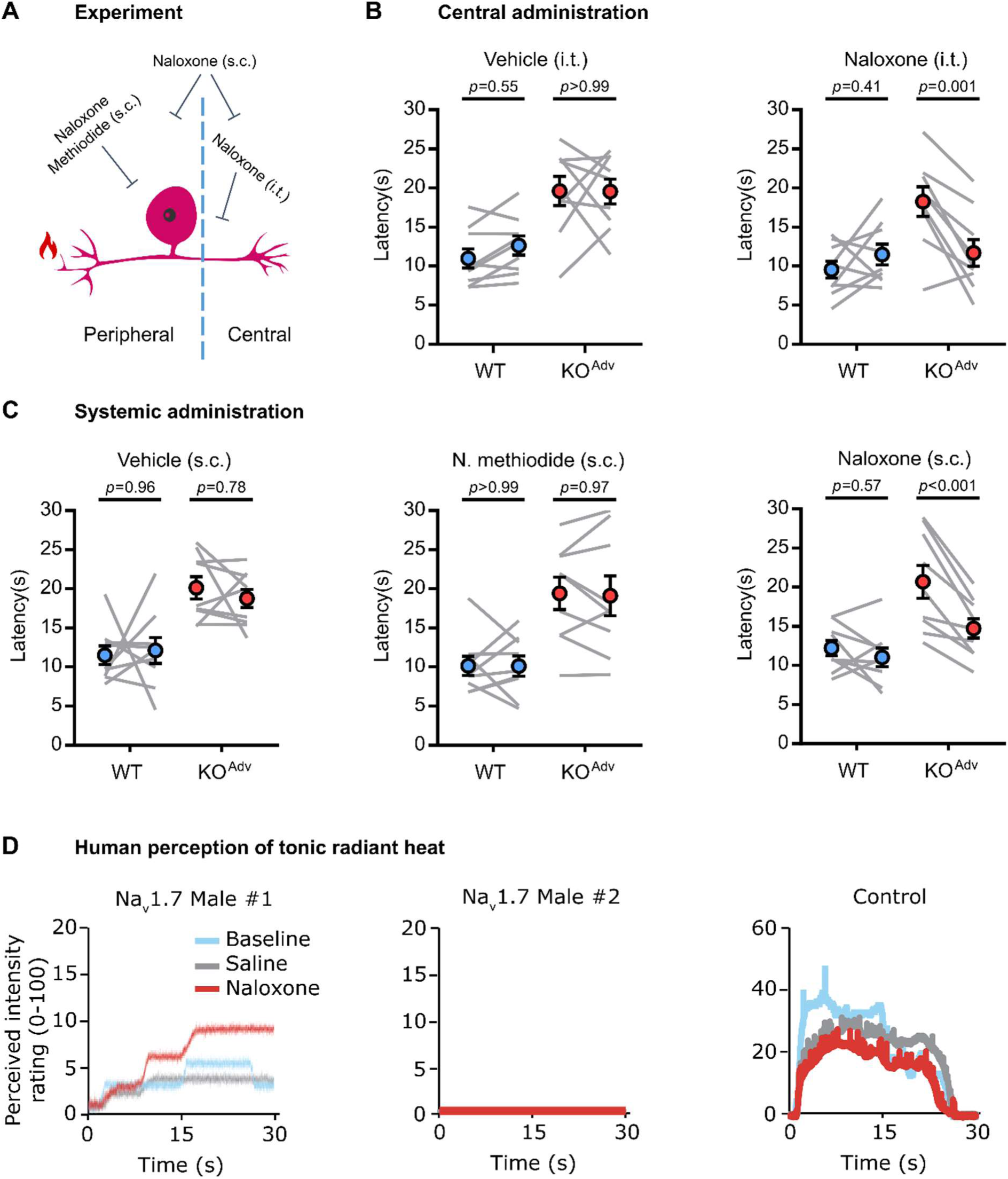
Blocking central opioid receptors reverses analgesia in mice and humans lacking Na_V_1.7. (A) Schematic of behavioural pharmacology experiment. (B) Behavioural assessment of the effect of vehicle and the opioid receptor blocker naloxone (3 mM in 5 µl for 20 minutes) administered centrally by intrathecal injection. (C) Behavioural assessment of the effect of vehicle and the opioid receptor blockers naloxone (2 mg/kg for 20 minutes) and naloxone methiodide (N. methiodide, 2 mg/kg for 20 minutes) administered systemically by subcutaneous injection. N. methiodide is peripherally-restricted and does not cross the blood brain barrier. (D) Line plots showing the reported, perceived intensity of tonic, radiant heat stimuli (45-48°C) in two newly-reported male Na_V_1.7 null individuals and one control participant, at baseline, during saline administration and after treatment with naloxone (12 mg). Naloxone appears to increase heat sensitivity in one Na_V_1.7 null participant (“male 1”), replicating previous observations in a single female null participant.^12^ Naloxone had no effect in a second Na_V_1.7 null participant (“male 2”). The control participant shows higher perceived pain intensity, which is not enhanced by naloxone. For both (B) and (C), the error bars represent standard error of the mean. Mean latencies before and after drug treatment were compared using repeated measures Two-Way ANOVA followed by post-hoc Sidak’s test. n=9 animals for WT and n=9 animals for KO^Adv^.

We have shown before that naloxone infusion restores nociception in a single rare female Na_V_1.7 null human.^12^ Here we extended these findings to two male humans with compound heterozygous Na_V_1.7 loss-of-function mutations (Table S1).^11,27,28^ We applied tonic radiant heat stimuli (25 s) to the forearm, while participants rated online the perceived intensity using a visual analogue scale (Figure 7D). In Male 1, naloxone strongly enhanced pain sensation, mimicking the effect on the previously reported female participant.^12^ In Male 2, there was no apparent effect of naloxone. In an age and gender-matched control, pain ratings were consistently higher than the Na_V_1.7 null individuals and not enhanced by naloxone. Thus, in 2 of the 3 human with Na_V_1.7 null mutations we have tested so far, block of opioid receptors enhances pain sensitivity, supporting a role for opioid signalling in driving analgesia associated with Na_V_1.7 loss-of-function.

## Discussion

Deletion of *SCN9A* encoding the peripheral sodium channel Na_V_1.7 in humans causes pain insensitivity and anosmia.^2,3^ Peripheral neuron Na_V_1.7 KO mice are pain-free in assays of mechanical, heat and inflammatory pain and show decreased spinal cord wide-dynamic range neuron firing to noxious stimuli.^15,29^ The main driver of analgesia must thus be the loss of Na_V_1.7 in sensory neurons. Because Na_V_1.7 exhibits slow closed-state inactivation, the channel is less likely to inactivate during slow sub-threshold depolarizations. Na_V_1.7 therefore mediates a ramp current that is hypothesized to amplify noxious stimulus-induced generator potentials to trigger action potential firing at nociceptor peripheral terminals.^30^ In patch-clamp studies of cultured mouse sensory neurons and human iPSC-derived nociceptors, under certain recording conditions Na_V_1.7 deletion impairs action potential firing upon depolarization of the soma.^11,31^ Recording of compound action potentials in KO^Adv^ mice also suggest some deficits in peripheral excitability.^32^ Hence the prevailing hypothesis – and rationale for developing peripherally-targeted inhibitors – is that analgesia after Na_V_1.7 loss-of-function arises from reduced excitability of the nociceptor peripheral terminal.

Using *in vivo* imaging and electrophysiology, we directly tested whether loss of Na_V_1.7 causes peripheral silencing of nociceptors in live mice. To our surprise, nociceptor excitability at the level of the DRG was largely unchanged, with normal levels of heat, polymodal and silent nociceptors, and normal calcium and spike response profiles. Nonetheless, we did observe reduced numbers of cells responding to noxious mechanical stimuli, consistent with earlier *in vitro* recordings that found ∼30% of Na_V_1.7-deficient putative nociceptors are electrically silenced in culture.^20^ In the main, however, action potentials propagate as far as the soma in Na_V_1.7-deleted sensory neurons, indicating the locus of analgesia is unlikely to lie at the peripheral terminal, explaining the failure of peripherally-targeted inhibitors of Na_V_1.7 to relieve pain.^33^

Studies of humans with loss-of-function mutations in Na_V_1.7 posit die back of peripheral nerves as a significant mechanism of analgesia.^11,34,35^ In three human Na_V_1.7 null individuals, microneurographic recording found evidence for Aδ but not C fibres, based on activity-dependent slowing profiles.^11^ These subjects were not challenged with painful stimuli. Nociceptors encompass all classes of sensory fibre including Aβ fibres and are defined by their ability to respond to and encode noxious stimuli.^22,23^ The preservation of neurogenic inflammation combined with a case report of a child with normal epidermal innervation and loss of pain are inconsistent with neuropathy as the principal mechanism of analgesia.^11,36^ Na_V_1.7 null individuals also lose the channel from innocuous touch sensing neurons that function normally and do not seem to die back.^37^

Na_V_1.7 is expressed along the length of sensory neurons, including at the central terminal, where it associates with many proteins including synaptotagmin-2.^7,25^ Given normal peripheral excitability, but reduced spinal cord neuron firing in Na_V_1.7 KOs, an alternative analgesic mechanism is that nociceptive input is lost through a failure of synaptic transmission from nociceptors to CNS neurons. Indeed, application of Na_V_1.7 inhibitors to spinal cord slices reduces synaptic transmission from afferents.^17,38^ Using glutamate imaging, electrophysiology and a substance P immunoassay, we observed deficits in pain-related neurotransmitter release in the spinal cord of mice lacking Na_V_1.7. As we directly activated dorsal roots of spinal cord slices, these experiments preclude any contribution of impaired peripheral excitability to the observed synaptic deficits. Echoing this, direct stimulation of dorsal roots in one CIP patient failed to elicit pain.^39^

How does loss of Na_V_1.7 impair neurotransmitter release? Analgesia in mice and humans lacking Na_V_1.7 can be substantially reversed by opioid antagonists or opioid receptor deletion. This results from both enhanced PENK production and opioid receptor signalling.^12–14^ Interestingly, regulation of both *Penk* transcription and opioid receptor signalling have been linked to lowered sodium levels that may contribute to Na_V_1.7 loss-of-function analgesia.^12,13^ As opioids are known to potently suppress substance P and glutamate release from spinal cord afferent terminals, we wondered if the synaptic impairments we saw in Na_V_1.7 KOs are dependent on opioid receptors.^40,41^ Peripheral excitability was unaffected by the opioid blocker naloxone in Na_V_1.7 KOs, consistent with previous findings using Na_V_1.7-deficient iPSC-derived human nociceptors.^11^ The deficits in both glutamate and substance P release were however reversed by naloxone. Both µ and δ opioid receptors are expressed at the central terminals in non-overlapping sets of nociceptors, explaining why both subtypes are required for analgesia.^14,42,43^

Anosmia in Na_V_1.7 nulls is wholly explained by impaired synaptic transmission from first-order olfactory sensory neurons, although somatic excitability to odorant stimuli is normal.^3^ We found that synaptic transmission from olfactory sensory neurons was not rescued by naloxone treatment. This is unsurprising given olfactory sensory neurons do not express opioid receptors.^44^ Na_V_1.7 is the only sodium channel available in olfactory sensory neuron nerve terminals; thus its loss completely blocks electrical activity pre-synaptically, consistent with the abolition of transmitter release by TTX.^3,45^ In contrast, nociceptors express other sodium channels that can support synaptic transmission in the absence of Na_V_1.7 when the inhibitory effect of opioids is removed.^46,47^

Opioid action at the central terminal is causally involved in pain insensitivity because central administration of naloxone was sufficient to substantially reverse analgesia, but systemic administration of a peripherally-restricted opioid antagonist was not. In humans, naloxone infusion enhanced sensitivity to nociceptive stimuli in 2 of the 3 Na_V_1.7 null individuals thus far tested. Why did naloxone have no effect on heat sensitivity in Male 2? This participant is hyposensitive to warmth, and during adolescence developed the ability to detect noxious stimuli by a ‘tingling’ sensation that was not perceived as unpleasant.^27^ These unusual phenotypic characteristics could impact his perception of the tonic heat stimulus used here, which he rated zero throughout. Importantly, numerous early case reports in the clinical literature attest to the dependence of CIP-like phenotypes on the opioid system, while sodium channel blockers show synergistic analgesia when paired with opioid drugs.^48–50^ Further studies support the role of CSF opioids within the HPA axis that can modulate spinal nociceptive processing.^51,52^ In one particularly elegant experiment, intrathecal injection of CSF from an unmapped CIP patient reduced heat nociception in rats. As this effect was blocked by naloxone, CSF opioids acting centrally are sufficient to recapitulate some CIP-associated analgesia in animals.^53^

What are the implications of a central, opioid-dependent mechanism of analgesia for pain therapies targeting Na_V_1.7? According to recent single-cell RNA sequencing studies, Na_V_1.7 is widely expressed in all sensory neuron subtypes, apart from proprioceptors.^37^ Yet mice and humans lacking Na_V_1.7 are insensitive only to noxious stimuli. If Na_V_1.7 deletion results in electrical silencing of sensory neurons, why is touch sensation not lost? Single-cell RNA sequencing data shows opioid receptors are not expressed by neurons expressing markers for low-threshold mechanoreceptors.^37^ We suggest that only in cells expressing opioid receptors can Na_V_1.7 deletion drive impaired synaptic transmission by enhancing opioid receptor signalling. Importantly, deletion of the transcription factor NFAT5 leads to elevated *Penk* mRNA transcripts, with no analgesia.^14^ Enhanced opioid receptor signalling is therefore a crucial component for analgesia in Na_V_1.7 KO mice. This is consistent with recent observations that peptide antagonists of Na_V_1.7 that elicit analgesia in mice can be inhibited by naloxone.^54,55^ Profound analgesic synergy between Na_V_1.7 blockers and low-dose opioids has also been reported.^8,38^ Na_V_1.7 block sensitizes opioid signalling only in nociceptors, reducing the effective concentration of opioids required to inhibit synaptic transmission from terminals. Close to 100% channel block may be required to drive the changes in opioid function responsible for analgesia.^12^ For intractable chronic pain, gene therapy strategies that mimic genetic loss-of-function will likely be required.^56^

Taken together, these data support a mechanism where pain insensitivity of mice and humans lacking Na_V_1.7 principally involves opioid receptors (Figure S6). Diminished peripheral excitability and die-back may also contribute to analgesia, but opioid-mediated suppression of neurotransmitter release plays a major role. Endogenous opioids inhibit synaptic communication between the central terminals of peripheral nociceptors and post-synaptic neurons in the spinal cord, resulting in diminished nociceptive input to the central nervous system. Dorsal horn neuron excitability is also diminished in Na_V_1.7 null mice.^57^ Importantly, nociceptor activity at the level of the dorsal root ganglion is largely unaffected by Na_V_1.7 deletion, despite behavioural analgesia. The critical locus of analgesia in Na_V_1.7 nulls is therefore the central terminal and not, as once thought, the periphery. Our findings consequently provide a biological explanation for the failure of peripherally-targeted Na_V_1.7 inhibitors to cause analgesia, and point to central terminal Na_V_1.7 and associated opioid signalling pathways as alternative therapeutic targets for pain relief.

## Acknowledgements

We thank the Wellcome Trust (200183/Z/15/Z) and the MRC. DIM was supported by a PhD fellowship from the Wolfson Foundation. SS is Versus Arthritis Fellow & Lecturer in Sensory Biology, and Versus Arthritis supported APL, QM and JNW with a programme grant (20200). FZ was supported by Deutsche Forschungsgemeinschaft grant Sonderforschungsbereich 894/A17. RMB was supported by Wellcome grant 110193, and Brain Research UK. We thank Xinzhong Dong for Pirt-GCaMP3 mice, Sonia Santana-Varela for help with mouse colonies, and Marco Beato and Filipe Nascimento and members of the Molecular Nociception Group for help and advice.

## Author contributions

The study was conceived by DIM, ECE and JNW. Imaging experiments were carried out by DIM, APL and ECE, electrophysiology by SS, APL, DIM and SRAA. Neurotransmitter release assays were carried out by DIM, QM and RMB. Olfaction studies were carried out by JW, MP and FZ. Behavioural assays were carried out by DIM, APL and QM. Human studies were carried out by FM and GDI. JZ and JJC provided reagents and advice. The paper was written by DIM and JNW with contributions from all authors.

## Declaration of interests

The authors declare no conflict of interest.

**Figure S1.**
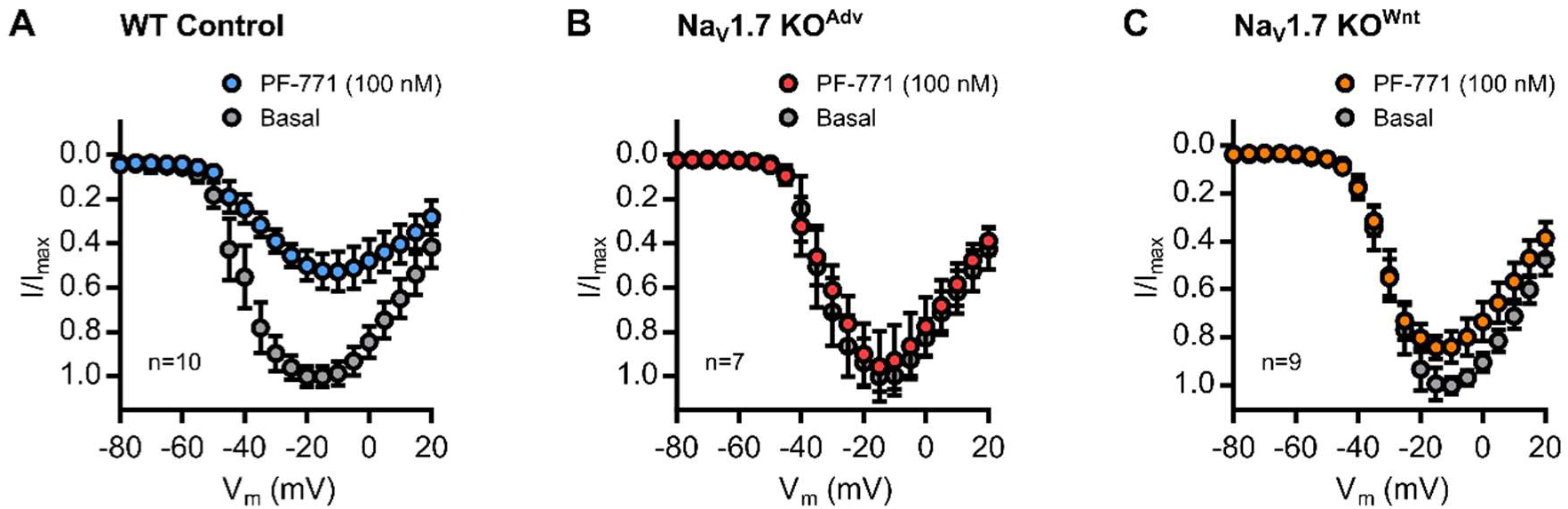
(*related to Figure 1*). Conditional deletion of *SCN9A* in sensory neurons abolishes voltage-gated sodium currents attributed to Na_V_1.7. (A) Current-voltage (I-V) curve showing that application of the Na_V_1.7-specific antagonist PF-05089771 (PF-771, 100 nM for 5 minutes) reduces voltage-gated sodium currents in WT sensory neurons. n=10 cells. (B) I-V curve showing PF-771 has no effect on voltage-gated sodium currents in sensory neurons from Na_V_1.7 KO^Adv^ mice. n=7 cells. (C) I-V curve showing PF-771 has no effect on voltage-gated sodium currents in sensory neurons from Na_V_1.7 KO^Wnt^ mice. n=9 cells. Error bars denote standard error of the mean.

**Figure S2.**
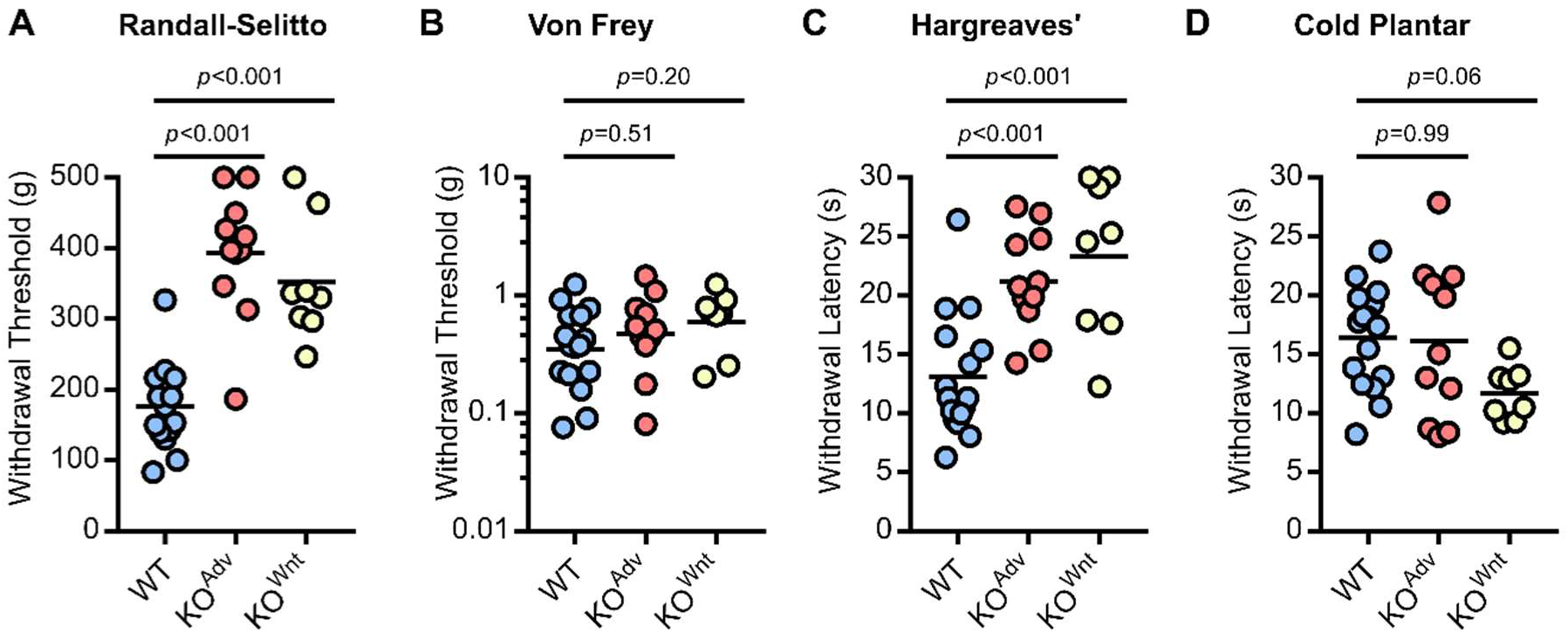
(*related to Figure 1*). Peripheral Na_V_1.7 knockout mice show reduced pain sensitivity. (A) Both Na_V_1.7 KO lines show increased tail withdrawal thresholds to noxious mechanical stimuli on the Randall-Selitto test. (B) Na_V_1.7 deletion has no effect on withdrawal thresholds to punctate tactile stimuli on the Von Frey test (C) Both Na_V_1.7 KO lines show increased withdrawal latencies to radiant heat stimuli on the Hargreaves’ test. (D) Na_V_1.7 KO^Adv^ show unchanged withdrawal latencies to dry ice stimuli on the Cold Plantar test compared to control, however Na_V_1.7 KO^Wnt^ display a small, but insignificant, hypersensitivity. For (A) to (D), means for each KO line were compared to WT control using One-Way ANOVA followed by post-hoc Dunnett’s test. n=16 mice for WT, n=11 mice for KO^Adv^ and N=8 mice for KO^Wnt^.

**Figure S3.**
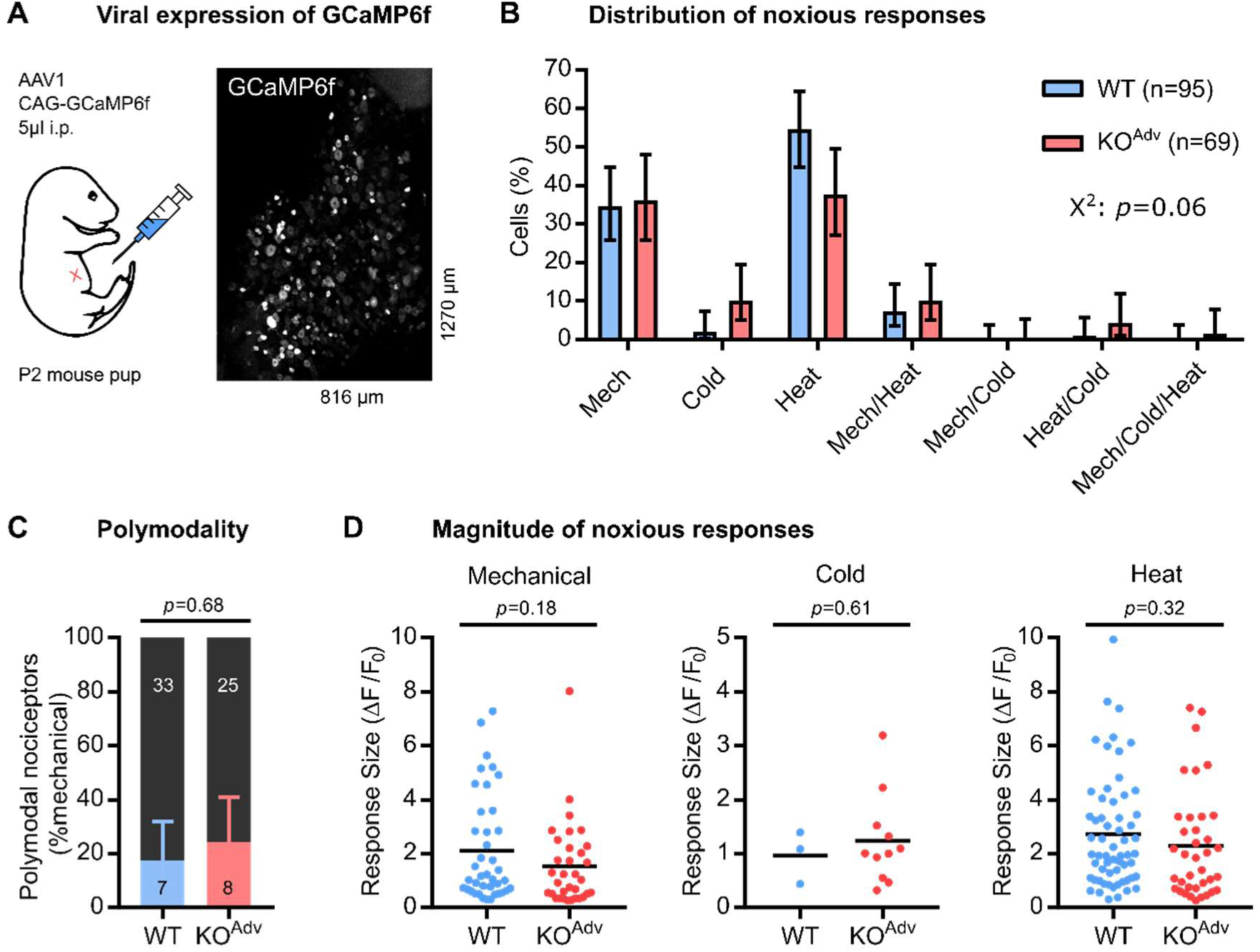
(*related to Figure 1*). *In vivo* calcium imaging of Na_V_1.7-deficient sensory neurons virally expressing GCaMP6f. (A) Confocal z-stack of DRG *in vivo* from a Na_V_1.7 KO^Adv^ mouse virally expressing GCaMP6f. AAV1-CAG-GCaMP6f was delivered by intraperitoneal injection into P2 mouse pups. (B) Bar plot summarizing the distribution of all sensory neurons that responded to different noxious stimuli in WT and KO^Adv^ animals. The error bars represent 95% confidence intervals and proportions were compared using Chi-Square test (χ^2^). n=95 cells from 4 WT mice (blue) and n=69 cells from 4 KO^Adv^ mice (red). Markedly fewer cells responded to cold in these animals compared to in Pirt-GCaMP3 mice, likely due to biased expression of the virally-delivered GCaMP6f. (D) Bar plot showing similar prevalence of polymodal nociceptors in WT and KO^Adv^ mice. Polymodal nociceptors are defined as pinch-sensitive neurons that respond to any noxious thermal stimulus (colour) and are expressed as a fraction of mechanically-sensitive cells (black). The error bars represent 95% confidence intervals and proportions were compared using the Chi-Square (χ^2^) test with Yates Correction. n=40 cells from WT and n=33 cells from KO^Adv^. (E-F) Scatter plots showing similar peak calcium responses (ΔF/F_0_) evoked by different noxious stimuli for WT and KO^Adv^. Mean response magnitude of KO^Adv^ was compared to WT control using an unpaired t test. Mechanical: n=40 cells from WT and n=33 cells from KO^Adv^. Cold: n=3 cells from WT and n=11 cells from KO^Adv^. Heat: n=60 cells from WT and n=37 cells from KO^Adv^.

**Figure S4.**
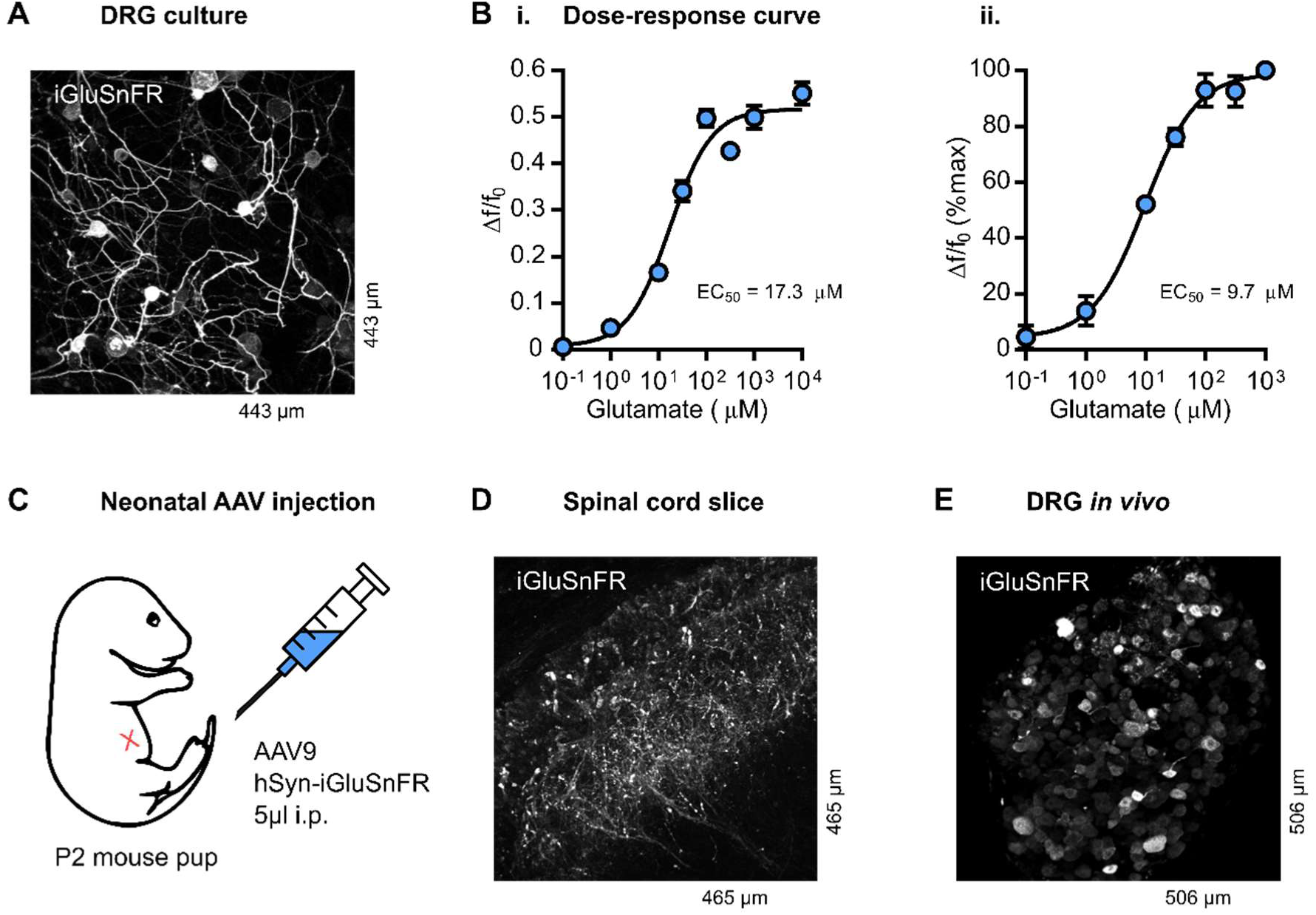
(*related to Figure 4*). Expression and function of iGluSnFR in sensory afferents. (A) Confocal z-stack of iGluSnFR-expressing cultured dorsal root ganglia neurons, *in vitro*. (B) Dose-response curve (i.) of iGluSnFR fluorescence (ΔF/F_0_) in cultured DRG neurons against extracellular glutamate concentration. n=36-154 neurons depending on tested concentration. EC_50_=17.3 µM. r^2^=0.426. Dose-response curve (ii.) of normalized iGluSnFR fluorescence (% maximum) in cultured DRG neurons against extracellular glutamate concentration. In this experiment, each cell was exposed to all glutamate concentrations tested. n=45 neurons. EC_50_=9.7 µM. r^2^=0.625. (C) Schematic of neonatal virus injection. (D) Two-photon z-stack of iGluSnFR-expressing sensory afferent terminals in dorsal horn of transverse spinal cord slice, *ex vivo*. (E) Confocal z-stack of iGluSnFR-expressing sensory afferent cell bodies in L4 dorsal root ganglion, *in vivo*.

**Figure S5.**
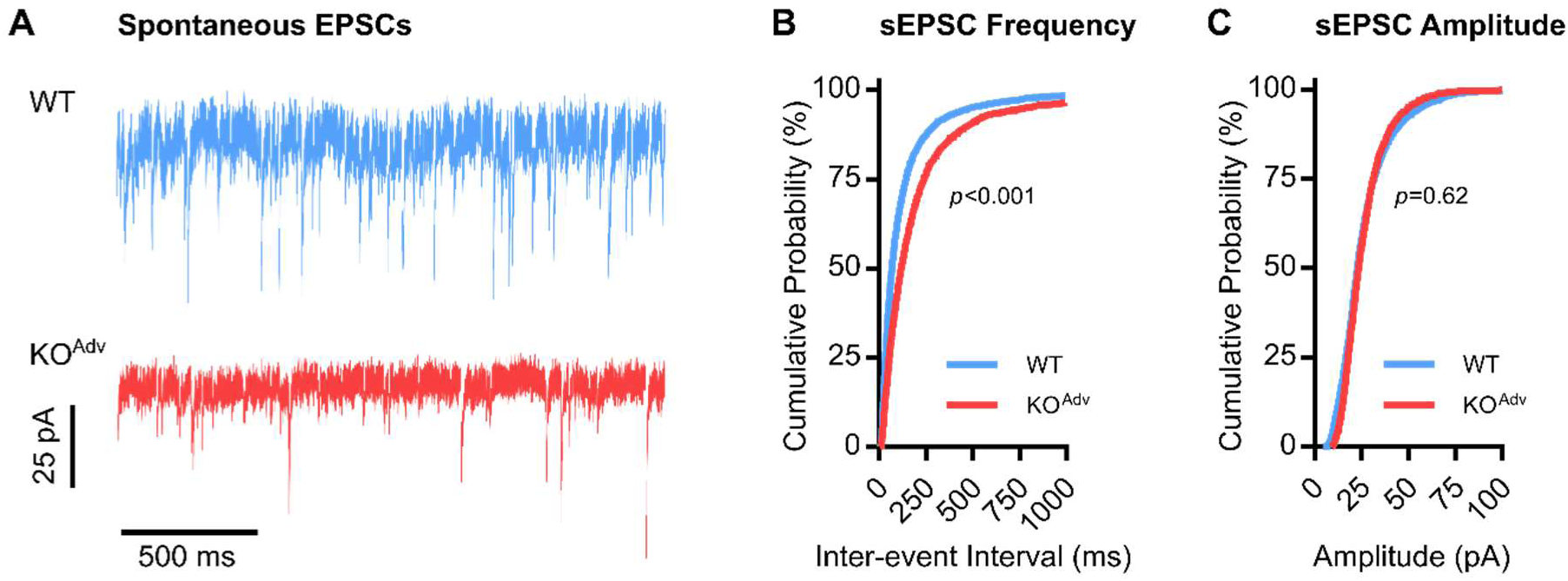
(*related to Figure 4*). Na_V_1.7 deletion reduces frequency, but not amplitude, of spontaneous excitatory post-synaptic currents. (A) Example traces showing spontaneous excitatory post-synaptic currents (sEPSCs) recorded from lamina II neurons in WT and KO^Adv^ mice. (B) Cumulative probability plots showing sEPSC frequency is reduced in KO^Adv^ animals. (C) Cumulative probability plots showing sEPSC amplitude is unaltered in KO^Adv^ animals. Means were compared using unpaired t tests. n=6972 events from 40 WT slices, and n=2406 events from 23 KO^Adv^ slices.

**Figure S6.**
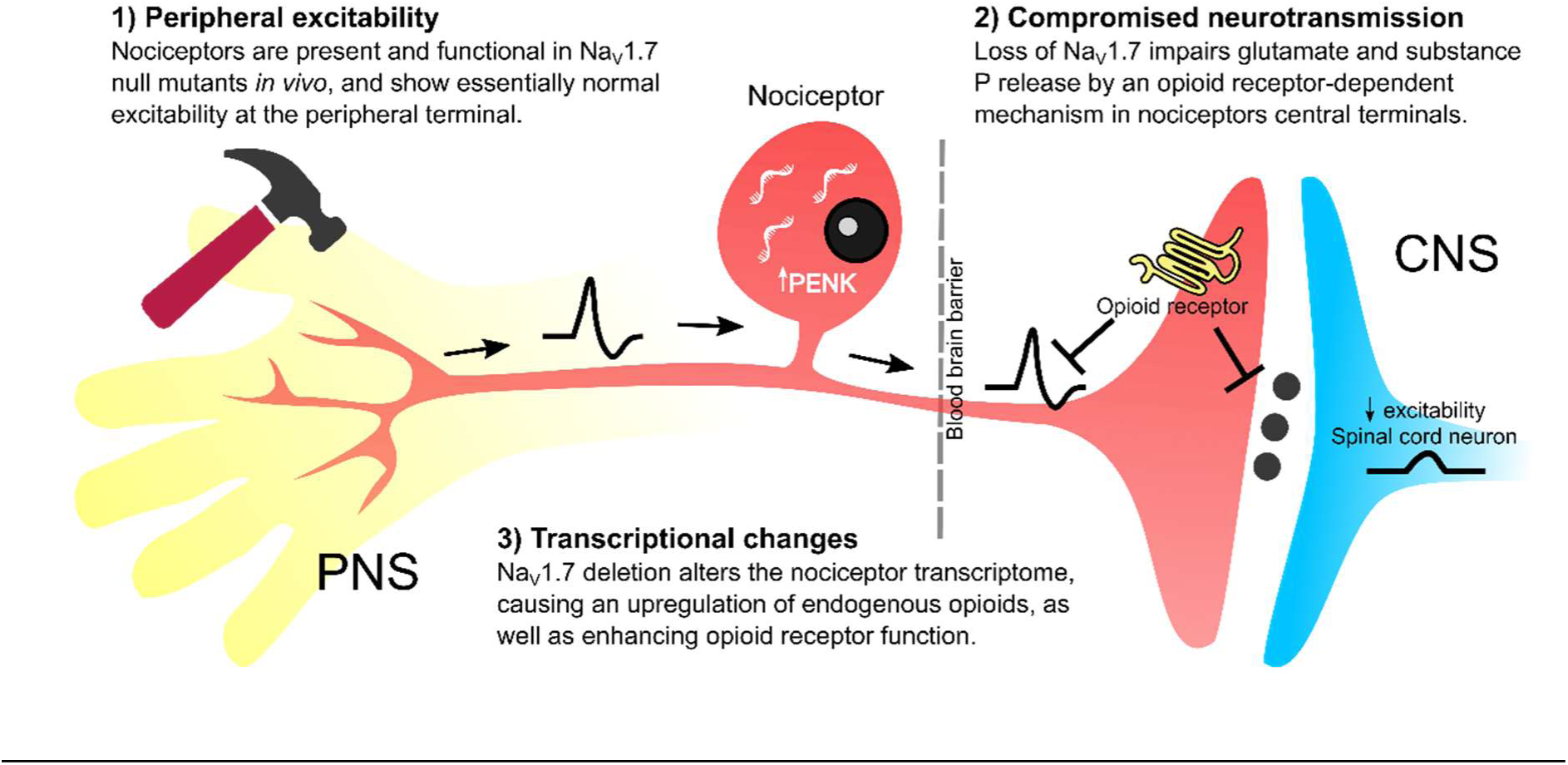
(*related to Figure 7*). Mechanisms of analgesia after loss of Na_V_1.7. Cartoon showing the effects of Na_V_1.7 deletion on nociceptor function. 1) *Peripheral excitability:* Some, but not all, humans with Na_V_1.7 null mutations show reduced intra-epidermal fibre density, while Na_V_1.7-deficient sensory neurons are less excitable *in vitro*. But, *in vivo*, terminal excitability is essentially normal, and neurogenic inflammation is preserved. Thus the peripheral terminal is not the locus of analgesia. 2) *Compromised neurotransmission:* Synaptic transfer from nociceptor terminals to spinal cord dorsal horn neurons is impaired after loss of Na_V_1.7. These synaptic deficits depend on opioid receptors, which can suppress both synaptic release and terminal excitability. Dorsal horn neurons are also less excitable due to absent post-synaptic Na_V_1.7. 3) *Transcriptional changes:* Loss of Na_V_1.7 leads to an upregulation of PENK, encoding pre-preproenkephalin, resulting in increased endogenous opioid signalling. Concomitantly, reduced sodium ingress following deletion of Na_V_1.7 leads to enhanced opioid receptor function inhibiting neurotransmitter release. Analgesia in mice and humans lacking Na_V_1.7 is thus dependent on opioids.

**Table S1.** (*related to Figure 7*). Mutations in *SCN9A* of human participants.

| PARTICIPANT | MUTATION | CHARACTERIZATION |
| --- | --- | --- |
| Male 1 | c.377+5C>T (intronic) – point mutation in splice donor site in intron 3 resulting in the use of a cryptic splice donor site, leading to a frameshift and premature stop codon in exon 4.<sup>11,28</sup> | Pathogenic.<sup>28</sup> |
|  | c.2686C>T (R896W) – amino acid change in exon 16.<sup>11</sup> | No Nav1.7 current when expressed in HEK293T cells.<sup>11</sup> |
| Male 2 | c.2488C>T (R830X) – premature stop codon in exon 16.<sup>27</sup> | No Nav1.7 current when expressed in HEK293T cells.<sup>11</sup> |
|  | c.5318delA (FS1773) – 1 bp deletion in exon 27, that induces a frameshift at position 1773 in the C terminal domain.<sup>27</sup> | 8-fold reduction in Nav1.7 current density versus WT when expressed in HEK293T cells.<sup>11</sup> |

**Table S2.**
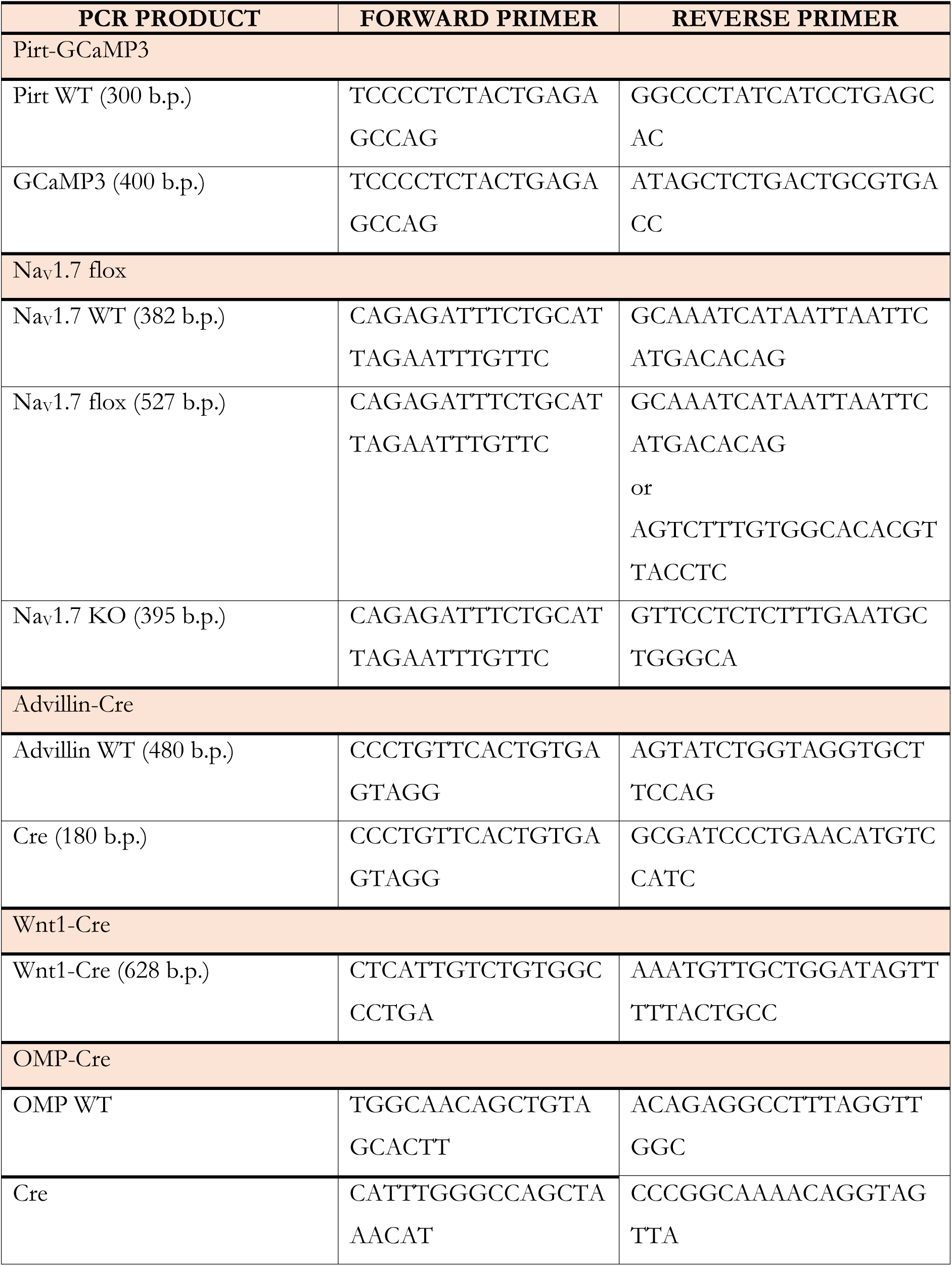
Primers used for mouse line genotyping.

## STAR* Methods

### Key Resources Table

**Experimental models: Organisms/Strains, Viruses, Pharmacological Agents**

| REAGENT or RESOURCE | SOURCE | IDENTIFIER |
| --- | --- | --- |
| Bacterial and Virus Strains |  |  |
| pAAV.CAG.GCaMP6f.WPRE.SV40 | Chen *et al*, 2013<sup>58</sup> | Addgene AAV1;<br>100836-AAV1 |
| pAAV.hSyn.iGluSnFr.WPRE.SV40 | Marvin *et al*, 2013<sup>24</sup> | Addgene AAV9;<br>98929-AAV9 |
| Chemicals, Peptides, and Recombinant Proteins |  |  |
| PF 05089771 | Tocris | 5931; CAS: 1430806-04-4 |
| Prostaglandin E2 | Merck (Sigma) | P0409; CAS: 363-24-6 |
| Naloxone hydrochloride dihydrate | Merck (Sigma) | N7758; CAS: 51481-60-8 |
| Naloxone Methiodide | Merck (Sigma) | N129; CAS: 93302-47-7 |
| Captopril | Merck (Sigma) | C4042; CAS: 62571-86-2 |
| Phosphoramidon disodium salt | Merck (Sigma) | R7385; CAS: 119942-99-3 |
| Dithiothreitol | Merck (Sigma) | D9779; CAS: 3483-12-3 |
| Thiorphan | Merck (Sigma) | T6031; CAS: 76721-89-6 |
| Bacitracin | Merck (Sigma) | B0125; CAS: 1405-87-4 |
| Experimental Models: Organisms/Strains |  |  |
| Mouse: Pirt-GCaMP3 | Kim *et al*, 2014<sup>59</sup> | N/A |
| Mouse: Nav1.7 flox | Nassar *et al*, 2004<sup>29</sup> | N/A |
| Mouse: Advillin Cre | Zhou *et al*, 2010<sup>60</sup> | N/A |
| Mouse: Wnt1 Cre | Danielian *et al*,<br>1998<sup>61</sup> | N.A |
| Mouse: OMP Cre Nav1.7 KO | Weiss *et al*, 2011<sup>3</sup> | N/A |
| Oligonucleotides |  |  |
| Primers for mouse genotyping | See Table S2 | N/A |
| Critical Commercial Assays |  |  |
| Substance P Parameter Assay Kit | R&D Systems | KGE007 |

| Software and Algorithms |  |  |
| --- | --- | --- |
| Fiji | ImageJ | https://fiji.sc/ |
| R studio | R | https://rstudio.com/ |
| LasX | Leica | https://www.leica-microsystems.com/ |
| PrairieView | Bruker | https://www.bruker.com/ |
| pCLAMP | Molecular Devices | https://www.moleculardevices.com/ |
| Prism | Graphpad | https://www.graphpad.com/ |

### Lead contact and materials availability

Please direct any requests for resources and reagents to the Lead Contact, John N. Wood. This study did not generate any new unique reagents.

### Experimental Model and Subject Details

#### Animals

All animal procedures carried out at University College London were approved by University College London ethical review committees and conformed to UK Home Office regulations. All animal procedures carried out at the University of Saarland were approved by the Institutional Animal Care and Use Committee of the University of Saarland (UdS) School of Medicine and were in accordance with the laws for animal experiments issued by the German Government.

The following mouse lines were used in this study: Advillin-Cre Na_V_1.7 KO, Wnt1-Cre Na_V_1.7 KO, Advillin-Cre Na_V_1.7 KO Pirt-GCaMP3, Wnt1-Cre Na_V_1.7 KO Pirt-GCaMP3 and OMP-Cre Na_V_1.7 KO. Breeding strategies were as previously described.^3,15^ Peripheral Na_V_1.7 knockout mice expressing GCaMP3 were generated by crossing knockout animals with mice homozygous for floxed Na_V_1.7 and homozygous for Pirt-GCaMP3.^59^ Mice were housed on a 12:12 hour light-dark cycle with food and water available *ad libitum*. For genotyping, genomic DNA was isolated from ear tissue or tail clip biopsy for PCR. Genotyping primers are summarized in Table S2. Both male and female animals were used for all experiments. The number of animals used to generate each dataset is described in individual figure legends.

#### Human subjects

We tested two male Na_V_1.7 null participants and one age and gender matched healthy control. All participants gave written informed consent. The study was approved by the UCL Research Ethics Committee. Both males are compound heterozygous nulls and have been previously described.^11,27^ Mutations are summarized in Table S2.

## Method Details

### Viral injections

Neonatal pups (P1-P3) were injected with 5 µl AAV1-CAG-GCaMP6f or AAV9-Synapsin-iGluSnFR via the intraperitoneal route using a Hamilton syringe connected to a 30G needle cannula. Care was taken to minimize exposure to foreign scents to ensure re-acceptance of pups by parents upon return to breeding cage.

### *In Vivo* Calcium Imaging

#### Acquisition

Mice expressing GCaMP3 or GCaMP6f (8 to 14 weeks, male and female) were anesthetized using ketamine (120 mg/kg) and medetomidine (1.2 mg/kg). Depth of anaesthesia was confirmed by pedal reflex and breathing rate. Animals were maintained at a constant body temperature of 37°C using a heated mat (VetTech). Lateral laminectomy was performed at spinal level L3-5. In brief, the skin was incised longitudinally, and the paravertebral muscles were cut to expose the vertebral column. Transverse and superior articular processes of the vertebra were removed using microdissection scissors and OmniDrill 35 (WPI). To obtain a clear image of the sensory neuron cell bodies in the ipsilateral dorsal root ganglion (DRG), the dura mater and the arachnoid membranes were carefully opened using microdissection forceps. The animal was mounted onto a custom-made clamp attached to the vertebral column (L1), rostral to the laminectomy. The trunk of the animal was slightly elevated to minimize interference caused by respiration. Artificial cerebrospinal fluid [containing 120 mM NaCl, 3 mM KCl, 1.1 mM CaCl_2_, 10 mM glucose, 0.6 mM NaH_2_PO_4_, 0.8 mM MgSO_4_, 18 mM NaHCO_3_ (pH 7.4) with NaOH] was perfused over the exposed DRG during the procedure to maintain tissue integrity, or the DRG was isolated by coating with silicone elastomer.

Images were acquired using a Leica SP8 confocal microscope (Leica). GCaMP3 or GCaMP6f was excited using a 488 nm laser (1-15% laser power). Images were acquired at 800Hz, with bidirectional laser scan. Typically, the pinhole was kept at 1 A.U, but in some experiments was increased to 1.5 A.U. to enhance brightness. Images magnification was between 0.75x and 3x optical zoom on a 10x air objective, depending on DRG anatomy. Noxious and innocuous stimuli were applied to the left hindpaw, ipsilateral to the exposed DRG. For thermal stimuli, the paw was immersed with ice-water (0°C) or water heated to 37°C or 55°C using a Pasteur pipette. For mechanical stimuli, we used noxious pinch with serrated forceps. PGE2 (500 µM in saline) was applied to the paw by intraplantar injection. Naloxone (2 mg/kg in saline) was delivered by subcutaneous injection into the scruff of the neck.

#### Analysis

Image stacks were registered to the first frame in the series using the FIJI plugin TurboReg (accurate rigid body transformation) to correct for XY drift. Stacks that showed excessive Z movement were excluded from analysis. Regions of interest (ROI) were manually drawn around apparently responding cells using the free hand tool in FIJI. Mean pixel intensity over time for each ROI was extracted and analysed. The time series of mean pixel intensity for each ROI was smoothened by a four time point moving average to remove high-frequency noise. Next, we calculated the derivative of the mean pixel intensity. We calculated a mean baseline derivative for the 10 s preceding stimulus application. Neurons were classed as responders if, within 30 s of stimulus application, the maximum derivative was greater than the baseline derivative plus five standard deviations – that is, a Z-score of at least 5. We then calculated the ΔF/F_0_ value for each response to obtain a normalized measure of change in fluorescence. Neurons which showed a ΔF/F_0_ less than 0.25 were then discarded. Each trace was then manually screened as a further precaution against false positives. The remaining neurons that made up the responding population were then used for statistical analysis.

### Behavioural Testing

All animal experiments were performed in accordance with Home Office Regulations. Observers were blinded to treatment and/or genotype. Animals were acclimatized to handling by the investigator and every effort was made to minimize stress during the testing. Both male and female animals were used.

#### Randall Selitto

The threshold for mechanonociception was assessed using the Randall Selitto test.^62^ Animals were restrained in a clear plastic tube. A 3 mm^2^ blunt probe was applied to the tail of the animal with increasing pressure until the mouse exhibited a nocifensive response, such as tail withdrawal. The pressure required to elicit nocifensive behavior was averaged across three trials. The cut-off was 500 g.

#### Von Frey

Punctate mechanical sensitivity was measured using the up-down method of Chaplan to obtain a 50% withdrawal threshold.^63^ Mice were habituated for one hour in darkened enclosures with a wire mesh floor. A 0.4 g Von Frey filament was applied to the plantar surface of the paw for 3 s. A positive response resulted in application of a filament of lesser strength on the following trial, and no response in application of a stronger filament. To calculate the 50% withdrawal threshold, five responses surrounding the 50% threshold were obtained after the first change in response. The pattern of responses was used to calculate the 50% threshold = (10[χ+κδ])/10,000), where χ is the log of the final von Frey filament used, κ = tabular value for the pattern of responses and δ the mean difference between filaments used in log units. The log of the 50% threshold was used to calculate summary and test statistics, in accordance with Weber’s Law.

#### Hargreaves’ Test

Spinal reflex responses to noxious heat stimulation were assessed using the Hargreaves’ test.^64^ Mice were habituated for an hour in plexiglass enclosures with a glass base. Before testing, the enclosures were cleaned of faeces and urine. Radiant heat was then locally applied to the plantar surface of the hindpaw until the animal exhibited a nocifensive withdrawal response. Average latencies were obtained from three trials per animal, with inter-trial interval of 15 mins. Cut-off time was 30 s. The effect of intraplanar PGE2 (500 µM) on heat sensitivity was assessed using the Hargreaves’ test. A baseline withdrawal latency was obtained and then measured again 10 minutes following PGE2 treatment. The effect of subcutaneous opioid blockers (2 mg/kg for 20 minutes) or intrathecal naloxone (3 mM in 5 µl) was also assessed using this assay, 20 minutes following drug treatment. For intrathecal injections, mice were anesthetized using 2-3% isofluorane and drugs delivered via a 30G needle cannula attached to a Hamilton syringe.

#### Cold Plantar

Spinal reflex responses to cooling were assessed using the Cold Plantar test.^65^ Mice were placed in Plexiglass enclosures with glass flooring and acclimatized for one hour. Before testing, faeces and urine were removed and the animal was left to settle. Dry ice was compacted into a blunt 2 ml syringe and applied to the glass surface just below the hindpaw. The time to withdrawal was measured. Cut off was 30 s. Testing was repeated 3 times, and averaged, with a waiting period of 15min between stimulations.

### Electrophysiology

#### In vitro electrophysiology

Mice were killed by inhalation of a rising CO_2_ concentration followed by cervical dislocation to confirm death. Dorsal root ganglia were dissected and then digested in an enzyme mix for 45 minutes before mechanical trituration. Neurons were re-suspended in DMEM supplemented with nerve growth factor and plated onto 12 mm glass coverslips coated with poly-L-lysine/laminin.

Patch pipettes (tip resistance of 3-5 MΩ) were filled with intracellular solution containing: 140 mM CsF, 1 mM EGTA, 5 mM NaCl and 10 mM HEPES. To isolate macroscopic sodium currents, neurons were continuously perfused with room temperature extracellular solution containing: 35 mM NaCl, 75 mM Choline-Cl, 30 mM TEA-Cl, 4 mM KCl, 1.8 mM CaCl_2_, 1 mM MgCl_2_, 10 mM HEPES, 5 mM Glucose and 0.1 mM CdCl_2_. Whole-cell recordings were obtained using an Axopatch 200B amplifier, filtered at 10 kHz and digitized at 50 kHz via a Digidata 1322A (Axon Instruments). Medium diameter neurons from WT and Na_V_1.7 KO mice were voltage-clamped at −70 mV. Series resistance compensation was at least 60%. To measure the voltage-dependence of sodium channel activation, the holding command was dropped to −120 mV to de-inactivate all sodium channels and then a step-protocol from −80 to 20 mV was applied, in increments of 5 mV, to activate sodium channels. To determine the contribution of Na_V_1.7 to the total sodium current, the Na_V_1.7 blocker PF 05089771 was applied for 5 minutes at 100 µM. As PF 05089771 is a state-dependent blocker that binds only to the inactivated state of the channel, the holding command was increased to −40 mV to inactivate sodium channels for the duration of drug application.

#### Ex vivo spinal cord slice electrophysiology

Spinal cord preparations were obtained from male or female mice, between 30 and 60 days old, from either wild-type C75Bl/6 (WT) or conditional Na_V_1.7 knockout (Na_V_1.7 KO). Animals were anesthetized via intraperitoneal injection of a ketamine/xylaxine mix (80 mg/kg and 10 mg/kg respectively) and decapitated. The spinal cord was dissected in ice cold aCSF containing: 113 mM NaCl, 3 mM KCl, 25 mM NaHCO3, 1 mM NaH2PO4, 2 mM CaCl2, 2 mM MgCl2, and 11 mM D-glucose. Once dissected free from the vertebral column, the spinal cord was carefully cleaned from connective tissues and dorsal roots were cut at approximately 2 mmm length. The spinal cord was then glued to an agar block and glued to the slicing chamber of a HM 650V vibratome (Microm, ThermoFisher Scientific, UK). The slicing solution contained: 130 mM K-gluconate, 15 mM KCl, 0.05 mM EGTA, 20 mM HEPES, 25 mM D-glucose, 3 mM kynurenic acid, 2 mM Na-Pyruvate, 3 mM Myo-Inositol, 1 mM Na-L-Ascorbate, and pH 7.4 with NaOH_4_. Slices were incubated for 40 minutes at 35 degrees and then allowed to equilibrate at room temperature for further 30 minutes before starting the recordings.

Voltage clamp recordings were performed using either a Molecular Devices Multiclamp 700B (Scientifica, UK) or an ELC-03X amplifier (NPI electronics, Germany). Signals were filtered at 5KHz, acquired at 50 KHz using a Molecular Devices 1440A A/D converter (Scientifica, UK) and recorded using Clampex 10 software (Molecular Devices, Scientifica, UK). Electrodes were pulled with a Flaming-Brown puller (P1000, Sutter Instruments, USA) from borosilicate thick glass (GC150F, Harvard Apparatus, UK). The resistance of the electrodes, following fire polishing of the tip, ranged between 3 and 5 MΩ. Bridge balance was applied to all recordings. Intracellular solution contained 125 mM K-gluconate, 6 mM KCl, 10 mM HEPES, 0.1 mM EGTA, 2 mM Mg-ATP, pH 7.3 with KOH, and osmolarity of 290–310 mOsm. Cells were targeted for patching in the inner and outer Lamina II and visualized through an Eclipse E600FN Nikon microscope (Nikon, Japan) equipped with infrared differential interference contrast (IR-DIC) connected to a digital camera (Nikon, DS-Qi1Mc). Cells were voltage-clamped at −70 mV and spontaneous excitatory post synaptic currents (sEPSCs) recorded. sEPSCs were automatically detected using ClampFit in a 5 s window for each cell.

### *Ex vivo* olfactory bulb electrophysiology

Acute MOB slices were prepared from 4 - 11 week old mice (male and female) anesthetized with CO_2_ before decapitation. OBs were rapidly dissected in ice-cold oxygenated (95% O_2_, 5% CO_2_) solution containing the following (in mM): 83 NaCl, 26.2 NaHCO3, 1 NaH2PO4, 2.5 KCl, 3.3 MgCl_2_, 0.5 CaCl_2_, 70 sucrose, pH 7.3 (osmolarity, 300 mOsm/l). The tissue was mounted on a vibratome (VT1000S; Leica Microsystems, Nussloch, Germany) and horizontal MOB slices (275 µm thick) were cut in the same solution. Slices were stored at 30 - 35° C for 15 - 20 min in standard extracellular solution and afterwards at room temperature until use. The extracellular solution contained the following (in mM): 125 NaCl, 25 NaHCO_3_, 2.5 KCl, 1.25 NaH_2_PO_4_, 1 MgCl_2_, 2 CaCl_2_ and 10 glucose (continuously bubbled with 95% O_2_, 5% CO_2_). Tissue slices were placed in the recording chamber and superfused at a rate of ∼2 ml/min (gravity flow) with extracellular solution bubbled with carbogen (95% O_2_, 5% CO_2_). Cells were visualized in intact tissue slices with a 40x water immersion objective lens (Olympus) using infrared-optimized differential interference contrast optics and fluorescent illumination and a GFP filter set attached to the microscope to elucidate the morphology of lucifer yellow-filled mitral and tufted cells (BX50WI, Olympus).

Slice recordings were carried out at room temperature using an EPC-9 automated patch-clamp amplifier (HEKA Elektronik, Lambrecht, Germany) and Pulse 8.11 software as described previously.^3^ Patch pipettes were pulled from borosilicate glass tubing (World Precision Instruments, Germany). The signals were filtered using an eight-pole Bessel filter built into the EPC-9 amplifier and digitized at a frequency ≥ filter cut-off frequency (VR-10B, Instrutech Corp.). The sampling rate during all recordings was 10 kHz. Recording pipettes had resistances of 3 - 6 MΩ. Cells were voltage-clamped in the whole-cell patch-clamp mode. M/T cells had an ellipsoid-shaped cell body with a diameter of >10 µm, were located in the mitral cell layer or in the external plexiform layer. We did not discriminate between mitral cells and tufted cells within the group of M/T cells. M/T cells were filled with lucifer yellow during the recording and were afterwards visually inspected using fluorescent illumination.

The intracellular solution contained (in mM): 140 CsCl, 1 EGTA, 10 HEPES, 2 ATP Na-salt, 1 GTP Mg-salt, 5 QX-314 (a lidocaine derivative; Sigma-Aldrich, Taufkirchen, Germany), 0.1 lucifer yellow, 0.4 neurobiotin (Vector Laboratories, Burlingame, CA, USA); pH 7.1; osmolarity 290 mosm). The theoretical liquid junction potential between intracellular and extracellular compartments was calculated to be 4.1 mV and was not corrected.

After establishing a whole-cell recording, cells were voltage clamped to −60 mV. We waited for at least 2 min before data acquisition began to allow for equilibration of intracellular solution into the dendrites. Electrical stimulation of the olfactory nerve layer was applied via a glass electrode filled with extracellular solution and connected to an Isolated Pulse Stimulator Model 2100 (A-M Systems Instruments, USA). Electrodes were visually positioned in close proximity to the corresponding glomerulus of the recording site and stimulus duration and intensity was 1 ms and 100 V, respectively. Extracellular solution containing the opioid receptor antagonist naloxone (300 µM, Sigma Aldrich, Germany) was perfused to the MOB slice for at least 10 minutes.

All electrophysiological data were analyzed using Igor Pro software (WaveMetrics) and Excel (Microsoft). For pharmacological experiments, amplitudes of evoked EPSCs were assessed. The Student’s t test was used to measure the significance of difference between two distributions. Data are expressed as means ± SEM.

### *In vivo* DRG electrophysiology

Electrophysiological recordings were performed by a blinded experimenter. Mice were anaesthetized with isofluorane (4%; 0.5 l/min N_2_O and 1.5 l/min O_2_) before being secured in a stereotaxic frame. Depth of anaesthesia was reduced and maintained at 1.5% isoflurane during the experiment. Lateral laminectomy was performed to expose the L4 DRG, as described above for *in vivo* imaging. Multi-unit extracellular recordings were made from DRG neurons using parylene-coated tungsten electrodes (A-M Systems). Mechanical and thermal stimuli were applied to the peripheral receptive field of hindpaw glabrous skin ipsilateral to the exposed DRG. Natural stimuli (dynamic brush, von Frey hairs 0.16–26 g, noxious prod 100 and 150 g/cm^2^ mechanical stimulation, thermal water jet 35–55°C and iced water) were applied in ascending order of intensity to receptive fields for 10 s and the total number of evoked spikes recorded. Ethyl chloride was applied for 1s as a noxious cold stimulus and the total number of evoked spikes in 10 s was quantified. Evoked activity of neurons was visualized on an oscilloscope and discriminated on spike amplitude and waveform basis using a CED 1401 interface coupled to Spike2 software (Cambridge Electronic Design) to record waveform templates and carry out principal component analysis. For naloxone experiments, 2 mg/kg naloxone in saline was injected subcutaneously into the scruff of the neck, and the stimulation protocol repeated again 20 minutes after naloxone injection.

### Glutamate Imaging

#### In vitro characterization

Dorsal root ganglia neurons were dissociated and cultured onto glass coverslips as described above. Neurons were treated with iGluSnFR AAV particles diluted in culture media at dilutions varying from 1/100 to 1/4000. After 2-5 days treatments, neurons expressed the virus at all tested dilutions. Coverslips were transferred to an imaging chamber and perfused with extracellular solution containing: 140 mM NaCl, 4 mM KCl, 1.8 mM CaCl2, 1 mM MgCl2, 10 mM HEPES and 5 mM glucose, with pH 7.4. Images were acquired using a Leica SP8 confocal microscope with 20x immersion objective and iGluSnFR was excited using a 488 nm laser (1-5% laser power). Glutamate was applied at various concentrations in the bathing solution resulting in fluorescence increases localized to the plasma membrane. For analysis, ring-shaped regions of interest were drawn around the membrane and mean pixel intensity extracted and converted to ΔF/F_0_. Three-parameter dose-response curves were fit using GraphPad prism with a standard Hill Slope of 1.

#### Two-photon imaging

Lumbar spinal cord slices were prepared for glutamate imaging from P9-P21 mice virally expressing iGluSnFr in sensory afferents.^24^ Mice were culled by intraperitoneal injection of a ketamine (60mg/kg) and xylazine (12mg/kg) cocktail followed by decapitation and exsanguination. Spinal cords were dissected in an ice-cold oxygenated (5% CO_2_ / 95% oxygen) dissection solution containing: 215 mM sucrose, 3 mM K-gluconate, 1.25 mM NaH_2_PO_4_, 26 mM NaHCO_3_, 4 mM MgSO_4_-7H_2_O, 10 mM d-glucose, 1 mM kynurenic acid and 1 mM CaCl_2_. Spinal cords were embedded in low-melting point agarose (2-3%) in ASCF and then sectioned using a vibrating microtome into 500 um thick transverse slices in oxygenated ACSF. Slices were incubated for at least an hour in oxygenated ACSF at 37°C. The ACSF contained: 111 mM NaCl, 3.085 mM KCl, 10.99 mM d-glucose, 25 mM NaHCO_3_, 1.26 mM MgSO_4_-7H_2_O, 2.52 mM CaCl_2_, and 1.1. mM KH_2_PO_4_.

Slices with dorsal roots attached were transferred to the recording chamber and pinned using a harp. Using a 10x air objective on a Brüker 2P microscope, the dorsal roots were visualized and approached with the suction electrode. Roots were gently suctioned into the suction electrode, allowing a tight seal to form. An Isoflex (Molecular Devices) stimulus isolator was used to deliver negative current pulses of varying amplitudes. The stimulus isolator was triggered directly from the imaging software (Prairie View) and stimulation was timelocked to image acquisition. The temporal profile of the output waveform was controlled from the imaging software, with pulse duration of 400 µs. To ensure accurate time-locking, the output from the stimulus isolator was also recorded by the imaging software.

Images were acquired using a 2-photon microscope (Bruker) with a 20x high NA water immersion objective. iGluSnFr was excited using a 920 nm laser line (Insight DS, Spectra-Physics) and 525/70 nm emission acquired by a GaAsP PMT (Hamamatsu) with gain set to maximum. The location of layer II of the dorsal horn was estimated using physical and fluorescent landmarks and a 250 x 125 pixel field of view drawn, equivalent to 195 x 98 µm. Images of glutamate release in response to different single pulse stimulus intensities were acquired at 10 Hz. Throughout the experiments slices were perfused with oxygenated room temperature ACSF and, in some experiments, the bathing solution contained naloxone (100 uM), which was applied for at least 20 minutes.

#### Analysis

Image stacks were analyzed in Fiji. Trials were concatenated and then registered to the first image in the stack (accurate rigid body transformation). Regions of interest were manually identified by a blinded experimenter and the mean pixel intensity over time per ROI extracted. Signals were converted to ΔF/F0. Glutamate release events time-locked to the stimulus were z-scored and considered significant if z>4. This dataset was then used for subsequent statistical analysis.

### Substance P ELISA

Thorico-lumbar transverse spinal cord slices were prepared from P14-40 WT and KO^Adv^ mice. Mice were culled by intraperitoneal injection of a ketamine (60mg/kg) and xylazine (12mg/kg) cocktail followed by decapitation and exsanguination. Spinal cords were removed from the spinal column by hydraulic extrusion and embedded in low-melting point agarose (2-3%) before sectioning using a vibratome (Leica) into 300 µm thick coronal slices in ice-cold oxygenated ACSF. Slices were then incubated for at least an hour in oxygenated ACSF at 37°C. The ACSF solution used throughout was the same recipe as used for glutamate imaging.

Following incubation, slices were treated with either 1 mM naloxone or vehicle for 20 minutes in 37°C oxygenated ACSF. Slices were then transferred to a 48 well plate, with 10 slices per well. To evoke substance P release, slices were treated with 2 uM capsaicin with and without naloxone for 20 minutes while being shaken at 300 rpm. The treatment solution also contained 0.1% BSA and a cocktail of peptidase inhibitors to maximize recovery of substance P. The cocktail contained (in µM): 200 captopril, 2 phosphoramidon disodium salt, 12 dithiothreitol, 20 thiorpan and 40 bacitracin. Following treatment, the superfusate was immediately collected and substance P concentration measured using a commercially-available Substance P competitive binding assay (R&D systems). All samples were run in duplicate (50 µl per sample) following the manufacturer’s instructions and optical density measured using a microplate reader. Sample substance P concentrations were interpolated from a standard curve run in parallel.

### Immunohistochemistry

One day prior to the experiment, mice were distributed onto individual homecages. Naloxone (cat# N7758, Sigma Aldrich) was dissolved in phosphate buffered saline (PBS) pH 7.4 and systemically applied by intraperitoneal injection of 2 mg/kg bodyweight (Pereira et al., 2018), while negative controls received vehicle (PBS) alone. Tested mice (n=3 each genotype) were Nav1.7^control^ [(flox/–)(OMP/–)(Cre/–)] and Nav1.7 KO^OMP^ [(fx/fx) (OMP/–)(Cre/–)] mice. For each mouse, three consecutive intraperitoneal injections with 30 min time intervals were performed. After each injection, mice were returned to their odour-rich home cages. At 24 hours after the last injection, mice were anesthetized, subjected to transcardial perfusion, followed by tissue preparation for tyrosine hydroxylase immunohistochemistry.

Mouse tissue preparation followed previously described methods.^3,67^ Adult mice (7-9 weeks-of-age) were anaesthetized using a mixture of 165 mg/kg body weight ketamine (Pharmacia GmbH, Berlin) and 11 mg/kg body weight xylazine (Bayer Health Care, Leverkusen), and were transcardially perfused with phosphate-buffered saline (PBS) pH 7.4, followed by 4% (w/v) paraformaldehyde in PBS. Olfactory bulbs (OBs) were dissected, incubated for 2 h in fixative and for overnight in 30% sucrose in PBS at 4°C, embedded in O.C.T. (Tissue-Tek), and snap-frozen in a dry ice/2-methylbutane bath. Frozen tissue sections (18 µm) were collected on a cryostat (Microm HM525, Walldorf, Germany) and stored at −80°C until subjected to immunohistochemistry. For tyrosine hydroxylase (TH) immunostaining, tissue sections were rinsed in PBS, treated for 1 h with blocking buffer containing 0.3% Triton X-100 and 4% normal horse serum (NHS, Vector Laboratories) prepared in PBS, followed by incubation in TH primary antibody (mouse monoclonal, cat# 22941, RRID:AB_572268, ImmunoStar, Hudson, WI, USA) diluted 1:2000 in blocking solution. Tissue sections were washed three times 10 min in PBS and incubated in Alexa-Fluor 488 conjugated goat-anti-mouse secondary antibody (1:1000, Thermo Fisher Scientific cat# A-11029, RRID:AB_2534088) for 1 h in the dark. Tissue sections were rinsed in PBS, the nuclei counterstained with Hoechst 33342 (1:10,000 in PBS, Invitrogen) for 10 min, rinsed again in PBS, and cover slipped using fluorescence mounting medium (DAKO). All procedures were conducted at room temperature with the exception of tissue incubation in primary antibody solution at 4°C.

Confocal fluorescence images were acquired on a Zeiss LSM 880 confocal microscope containing a 32-channel GaAsP-PMT and 2-channel PMT QUASAR detector. Each image shown in Figure 6C refers to a single 2.5 µm thick optical section. Images were assembled and minimally adjusted in brightness using Adobe PhotoShop Elements 10.

### Human Sensory Testing

Perception of tonic radiant heat was assessed at baseline and during intravenous administration of saline or naloxone (12 mg), in a randomized order. The perception of long-lasting, tonic nociceptive stimuli is generally considered to be less confounded by attention than transient noxious stimuli, involving rapid, attentional shifts that can confound perceptual measures.^68^ Psychophysical assessment was carried out by an experimenter blind to the pharmacological condition. Tonic radiant heat was generated by a CO_2_ laser, whose power is regulated using a feedback control based on an online measurement of skin temperature at the site of stimulation (Laser Stimulation Device, SIFEC, Belgium). The CO_2_ laser selectively stimulates both A-delta and C fibers. On each trial, tonic radiant heat was delivered to the forearm for 25 s and kept constant at either 45 or 48 °C.^69^ Participants were asked to rate the intensity of the thermal sensation on a visual analogue scale throughout the trial (0=no sensation, 100=worst pain imaginable). Three trials per stimulus temperature were given on each session (baseline, saline and naloxone) in a randomized order.

### Quantification and Statistical Analysis

For *in vivo* imaging experiments, n refers to the number of cells responding to any stimulus. For electrophysiology experiments, n refers to the number of recorded cells. For glutamate imaging experiments, n refers to the number of regions of interest. For all imaging and physiology data, the number of animals used is indicated in the legend. For behavioural and substance P experiments, n refers to the number of animals.

Datasets are presented using appropriate summary statistics as indicated in the legend. Error bars denote mean ± 95% confidence interval or mean ± SEM, as indicated in the legend. The 95% confidence interval around proportions was estimated using the Wilson-Brown method. Tests of statistical comparison for each dataset are described in detail in figure legends. For grouped data, we made the appropriate correction for multiple comparisons. We set an α-value of *p*=0.05 for significance testing and report all p-values resulting from planned hypothesis testing.

No sample size calculation was performed, however our samples are similar to those used in the field.

### Data and code availability

Data are available from the lead contact on reasonable request.

## Notes

### Competing Interest Statement

The authors have declared no competing interest.

